# Cryo-EM and Solid State NMR Together Provide a More Comprehensive Structural Investigation of Protein Fibrils

**DOI:** 10.1101/2024.05.30.596698

**Authors:** Blake D. Fonda, Masato Kato, Yang Li, Dylan T. Murray

## Abstract

The Tropomyosin 1 isoform I/C C-terminal domain (Tm1-LC) fibril structure is studied jointly with cryogenic electron microscopy (cryo-EM) and solid state nuclear magnetic resonance (NMR). This study demonstrates the complementary nature of these two structural biology techniques. Chemical shift assignments from solid state NMR are used to determine the secondary structure at the level of individual amino acids, which is faithfully seen in cryo-EM reconstructions. Additionally, solid state NMR demonstrates that the region not observed in the reconstructed cryo-EM density is primarily in a highly mobile random coil conformation rather than adopting multiple rigid conformations. Overall, this study illustrates the benefit of investigations combining cryo-EM and solid state NMR to investigate protein fibril structure.

**Significance:** The use of multiple techniques to structurally characterize proteins provides models that accurately describe molecular conformations better than a technique used in isolation. Combination approaches allow for the study of proteins not only as rigid objects, but rather dynamic molecules that “breathe” over time. Cryogenic electron microscopy and solid state nuclear magnetic resonance are used jointly to provide a more detailed model of the same protein fibrils, and each technique provides novel insights.

## Introduction

No single structural biology tool is capable of routinely characterizing all aspects of protein fibrillization, since the phenomena, both pathological and functional, is influenced by both rigid structure and conformational flexibility. The structural characterization of protein fibrillization has led the discovery that, for some neurodegenerative diseases, fibril structure is indicative of disease pathology. For instance, in amyotrophic lateral sclerosis and frontal temporal dementia, distinct fibril structures, or polymorphs, of the protein TDP-43 are associated with disease pathology (Arseni et al., 2021, Arseni et al., 2023). Beyond the importance of pathogenic protein fibrillization, ample work has shown that protein fibrils can be functional. For example, proper biofilm formation and the formation of long term memory are linked to protein fibrillization processes (Oeen et al., 2019). Importantly, non-rigid regions outside the immobilized core region of pathogenic and functional protein fibrils play a key role in their formation (Nagaraj et al., 2020, Ulamec et al., 2020).

Structural determination of amyloid-like protein fibrils has increasingly focused on the use of cryogenic electron microscopy (cryo-EM) to determine high resolution structures. Indeed, cryo-EM has recently been used to solve the structure of 76 different Tau protein fibril structures, and recent reviews discussing high throughput and relevant structure elucidation have been published (Lövestam et al., 2022a, Lövestam et al., 2022b; Scheres, 2020).

Despite being incredibly powerful at determining high resolution structures of the rigid fibril core region, cryo-EM characterizations of the more mobile and flexible regions outside of the core are much less prevalent. In addition, significant time must be spent on the grid screening to obtain well-behaved fibrils on the sample grid with optimal ice thickness. The application of proteases and detergents is often required to achieve fibril separation for robust fibril particle picking (Arseni, 2021). Furthermore, the fibril structure must have a visible twist to yield sufficient orientations for cryo-EM helical reconstruction. Therefore, protein fibril reconstruction with cryo-EM is limited to fibrils with amenable characteristics.

Solid state nuclear magnetic resonance (NMR) is able to characterize both rigid and highly mobile regions of fibrils that may exist in distinct conformations (Bax et al., 2013). Solid state NMR, unlike cryo-EM, does not require fibril separation. In addition, solid state NMR methods can characterize fibril structures without helical twist. Nevertheless, like cryo-EM, solid state NMR has its own set of challenges. For example, it often requires multiple samples with different isotopic labeling schemes; computational approaches exist for chemical shift resonance assignment, but the distance constraints required for high-resolution structures are often done manually and need a variety of different NMR experiments, some having low signal to noise ratios (Murray et al., 2017; Tuttle et al., 2016; Tycko, 2011).

The benefits of using complementary solid state NMR and cryo-EM techniques for studying protein fibrils have been previously discussed (Li et al., 2020). Publications comparing the results from the two techniques, but with differing sample preparations, have been performed for fibrils related to dementia like serum amyloid A, Tau, and β-amyloid, among other proteins (Mammeri et al., 2024; Qiang et al., 2012; Sundaria et al., 2022; Zhang et al., 2023). Reviews comparing the use of solid state NMR and cryo-EM for structural studies of protein fibrils have been published (Eisenberg et al., 2017; Iadanza et al., 2018; Li, 2020; Zadorozhnyi et al., 2024).

The purpose of this study, rather than a high-level review, is to use cryo-EM and solid state NMR on the *same fibril preparations* and therefore gain a better understanding of how the two techniques work in synergy to describe a particular protein fibril conformation. In this work, solid state NMR and cryo-EM characterizations of fibrils formed by the C-terminal domain of Drosophila Tropomyosin isoform I/C (Tm1-LC) demonstrate great agreement. Secondary structures observed in the high resolution cryo-EM reconstruction are faithfully reproduced by secondary structures predicted from solid state NMR chemical shift measurements. In addition, while the fuzzy coat on the exterior of the rigid fibril core is invisible in the cryo-EM density map, solid state NMR characterizes the residue-specific motions for these residues.

## Results

Recombinant Tm1-LC readily fibrilizes in a neutral pH buffer at a concentration of 5 mg/mL followed by sonification and incubation for a week with intermittent agitation at 4 °C. Negative stain electron micrographs in Figure 1a–b show well-dispersed fibrils. Similar fibril formation has been observed for His-tagged Tm1-LC (Sysoev et al., 2020). The amino acid sequence of Tm1-LC shown in Figure 1c reveals a dominance of polar Asn, Ser, and Thr residues and a lack of aromatic amino acids. The protein construct used here contains a non-native N-terminal Ser-Tyr tag for UV quantification of the protein concentration and is referred to as Tm1-LC for the remainder of this manuscript. Figure 1d and Supplementary Figure 1 show that common aggregation and fibril formation prediction programs do not strongly predict an extended region of fibril formation (Conchillo-Solé et al., 2007; Fernandez-Escamilla et al., 2004; Maurer-Stroh et al., 2010; Walsh, 2014).

**Figure 1.**
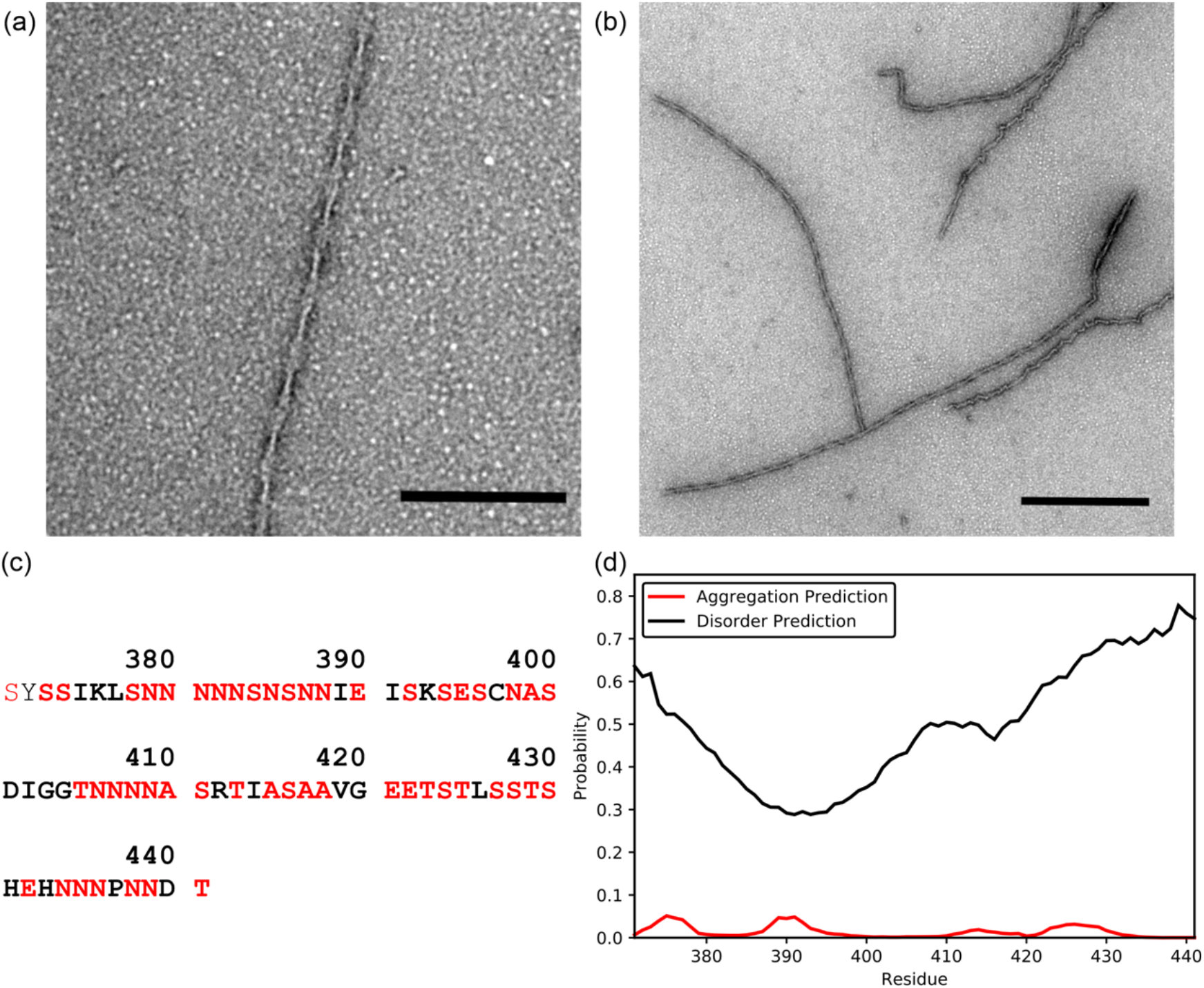
Negative Stain TEM of Fibrils and Predictions for the Tm1-LC Tail Domain. (a) and (b) Negative stain TEM micrographs of the Tm1-LC fibrils. The scale bar measures 200 nm. (c) The sequence from the wild-type Tm1-LC domain is in bold and the numbering scheme is relative to the full-length protein. The N-terminal Ser-Tyr tag used for UV quantification is unbolded. Dominant amino acids Ser, Asn, Ala, Thr, and Glu are in red. These five amino acids account for 70% of the sequence. (d) The PASTA 2.0 prediction of disorder and aggregation probability for Tm1-LC.

To obtain high-resolution details of the Tm1-LC fibril structure, cryo-EM data of Tm1-LC fibrils was collected and processed with RELION (He et al., 2017; Scheres, 2012). The 2D classes show distinct fibril conformations on the same grid (Supplementary Figure 2). The cryo-EM density maps in Figure 2a show the presence of four distinct fibril structures, three of which (classes A–C) are of sufficient quality for model building and/or refinement. Tables 1–2 and Supplementary Tables 1–2 detail the parameters of data collection, data processing, and model building. Supplementary Figure 3 depicts additional data for class A and provides additional analysis of the structure.

**Figure 2.**
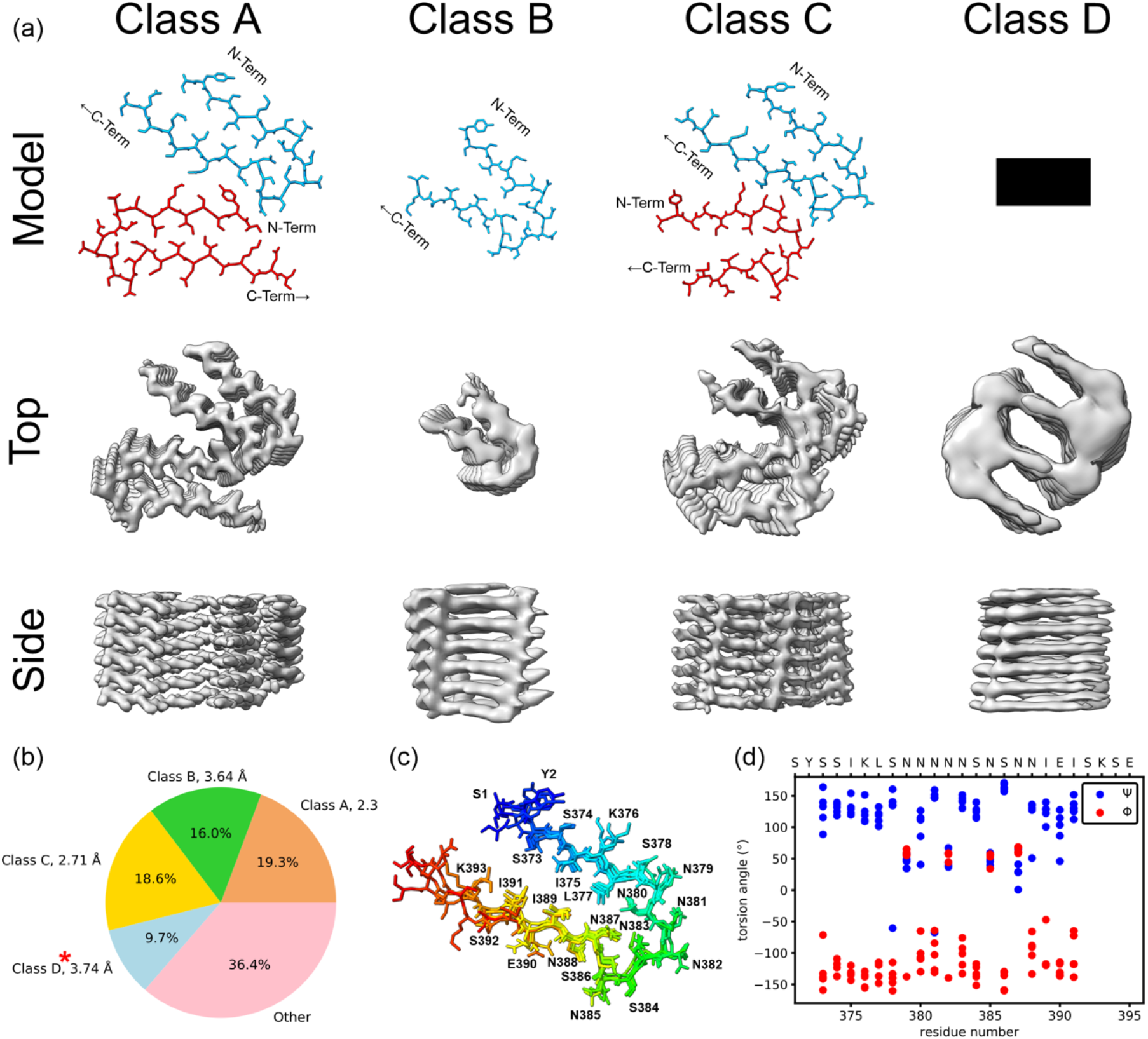
Cryo-EM Reconstruction of Tm1-LC Fibrils. (a) Model: structural models for the repeating unit of each fibril class. The black bar indicates no model could be built due to low resolution in the density map. Top: view of the cryo-EM density maps down the helical axis. Side: view of each cryo-EM electron density map perpendicular to the fibril long axis. (b) Percentage of the total picked particles for each class. See also Supplementary Figure 2. “Other” particles do not have consensus features. The resolutions noted in the labels are reported by RELION postprocessing. The red asterisk denotes a likely overestimate of resolution for class D due to a lack of secondary structure features. (c) Alignment of residues Y-S373–S392 for all five monomer models from classes A, B, and C. The N-terminal SY-tag is in dark blue. (d) Torsion angles extracted from the cryo-EM models. Blue represents the ψ angles and red represents the φ angles for each model of the Tm1-LC monomer. Torsion angles are plo_ed for residues that are structured across all classes. Supplementary Figure 5 shows the torsion angle plots by model class for all modeled residues.

**Table 1.** Cryo-EM data collection and reconstruction parameters. See methods for more information.

|  |  |  |  |  |
| --- | --- | --- | --- | --- |
| Microscope | Titan Krios (Thermo Fisher Scientific) |  |  |  |
| Camera | K3 (Gatan) |  |  |  |
| Acceleration voltage (kV) | 300 |  |  |  |
| Magnification | 105,000x |  |  |  |
| Defocus range ($\mu\text{m}$) | −1.0 to −2.4 | | | |
| Dose rate ($\text{e}^-/\text{\AA}/\text{s}$) | 8 | | | |
| Number of movie frames | 50 |  |  |  |
| Exposure time (s) | 4.5 |  |  |  |
| Total electron dose ($\text{e}^-/\text{\AA}$) | 52 | | | |
| Pixel size ($\text{\AA}$) | 0.83 | | | |
| Box size (pixel) | 320 |  |  |  |
| Inter box distance ($\text{\AA}$) | 14.25 | | | |
| Number of extracted segments | 2,295,590 |  |  |  |
| **Class Label** | **Class A** | **Class B** | **Class C** | **Class D** |
| Number of segments after 2D classification | 425,890 | 352,966 | 409,410 | 213,704 |
| Number of segments after 3D classification | 17,145 | 123,489 | 167,560 | 43,529 |
| Resolution, 0.143 FSC criterion ($\text{\AA}$) | 2.31 | 3.64 | 2.71 | 3.74 |
| Map sharpening B-Factor ($\text{\AA}^2$) | −28.84 | −100.33 | −77.96 | −82.96 |
| Helical rise ($\text{\AA}$) | 4.69 | 4.75 | 4.76 | 4.84 |
| Helical twist ($^\circ$) | +2.58 | +6.09 | +3.48 | +4.34 |

**Table 2.** Modeling Parameters for Class A. See methods for more information. Supplementary Tables 1 and 2 contain the statistics for classes B and C.

|  |  |
| --- | --- |
| Non-hydrogen atoms | 2376 |
| Number of protein chains | 12 |
| Map CC (around atoms) | 0.90 |
| RMSZ bonds (Å) | 0.004 |
| RMSZ angles (°) | 0.475 |
| All-atom clash score | 2.84 |
| Ramachandran favored/allowed/outliers (%) | 99.65/0.35/0 |
| Rotamer outliers (%) | 0 |
| CB outliers (%) | 0 |
| Molprobity score | 1.07 |

All four fibril 3D classes display typical cross-β fibril morphology with a helical repeat consistent with β-strand spacing at ∼4.75 Å. Additionally, all classes show a U-shaped fold consisting of two β-strands per Tm1-LC molecule that are separated by a brief, Asn-rich turn region. Classes A, C, and D show two molecules per repeating fibril unit, whereas class B has a monomeric repeating unit. These classes have relatively similar numbers of picked particles (Figure 2b; 10–20% of the total particles), although these percentages should not be taken as an exact measure of the ratio of conformations in the sample.

Alignment of the five separately modeled Tm1-LC monomer chains from classes A–C over residues Y-S373–S392 yields a backbone non-hydrogen atom RMSD of less than 1.5 Å, and a sidechain and backbone non-hydrogen atom RMSD of less than 2.1 Å, where Y-S373–S392 represents the continuous sequence from the Tyr in the N-terminal UV purification tag through residue S392. These small RMSD values show that the individual Tm1-LC molecules adopt the same secondary and tertiary structure, yet each fibril class has a unique quaternary structure. Additional RMSD values calculated from comparing the models are shown in Supplementary Tables 3–5. Alignment of all the models from classes A–C (Figure 2c) shows excellent agreement from the N-terminus until K393. For class D, the rigid body fitting of the class B monomer model into the class D density map shown in Supplementary Figure 4 indicates that the class D monomer conformation is similar to those from classes A–C. To further characterize the similarity of the fold across the modeled fibril classes, torsion angles for residues 373–391 are extracted and plotted in Figure 2d (see also Supplementary Figure 5). The small variation in torsion angles across the models, especially for β-strand regions S373–S378 and N388–I391, show a consistent conformation of Tm1-LC.

The primary difference between the Tm1-LC conformations lies in the quaternary structure packing interface, although it is not possible from the data to completely rule out an alternative fibril core residues for class D fibrils. Class D is consistent with an extended β-strand packing interface between both molecules of the repeating helical unit. Class A contains a packing interface with the N-terminus of one Tm1-LC molecule making intermolecular contacts with residues N385–E390 in the opposing molecule. Class C has a packing interface of N385–S392 making intermolecular contacts with K376–N382 on the second molecule. However, as discussed previously based on RMSD and torsion angle values, unique intermolecular contacts in classes A and C do not perturb the propensity of Tm1-LC to form the same tertiary structure. As shown for class A in Supplementary Figure 3, the Tm1-LC consensus fold across all classes has a hydrophobic interior and a packing interface/solvent-exposed exterior containing a mixture of polar and charged residues.

The models with the most residues in the rigid core are one monomer from class C and both monomers from class A which consist of residues SY-S373–S396. The consensus rigid core region across classes A, B, and C contains residues Y-S373–S392. Residues C397–T441 are not observed in all classes. Residues located downstream of the fibril core must therefore either adopt multiple conformations or are highly mobile and disordered.

The regions flanking the fibril core are investigated with solid state NMR experiments. Figure 3a shows the aliphatic region of an INEPT-TOBSY experiment recorded on ^13^C and ^15^N isotopically labeled Tm1-LC fibrils prepared identically to the sample used for the cryo-EM reconstruction. Scalar-based solid state NMR INEPT-TOBSY measurements report on the highly mobile regions of Tm1-LC in the fibrils (Andronesi et al., 2005; Hardy et al., 2001). The INEPT-TOBSY experiment selects regions of the protein with µs or shorter rotational correlation times (see methods for details). Manually determined sequence-specific non-ambiguous and ambiguous resonance assignments are overlayed on the spectrum in Figure 3a. Residues R412, I414, V419, G420, L426, N436, P437, and D440 are assigned unambiguously in the INEPT-TOBSY spectrum and lie outside of the region observed in the cryo-EM density maps. Figure 4a shows that the assigned residues from the INEPT-TOBSY spectrum have secondary chemical shifts nearly identical to their predicted random coil shifts (Kjaergaard et al., 2011; Kjaergaard, 2011; Maltsev et al., 2012; Schwarzinger et al., 2001; Wishart, 2011). Small secondary chemical shifts are interpreted as amino acid residues in a coil-like arrangement, i.e. non-β strand and non-α helical. Ambiguous assignments for additional signals in Figure 3a have chemical shifts consistent with intrinsically disordered Ala, Ser, Glu, and Asn residues. The tentative assignments for these resonances are depicted in red in Figure 3a and are also colored red in the secondary chemical shift plot in Figure 4a. Combination of the INEPT-TOBSY solid state NMR result with the cryo-EM reconstruction structurally reveals that the Tm1-LC fibrils consist of an N-terminal rigid core with a C-terminal flanking region that contains residues that retain significant motion despite being close to the rigid core.

**Figure 3.**
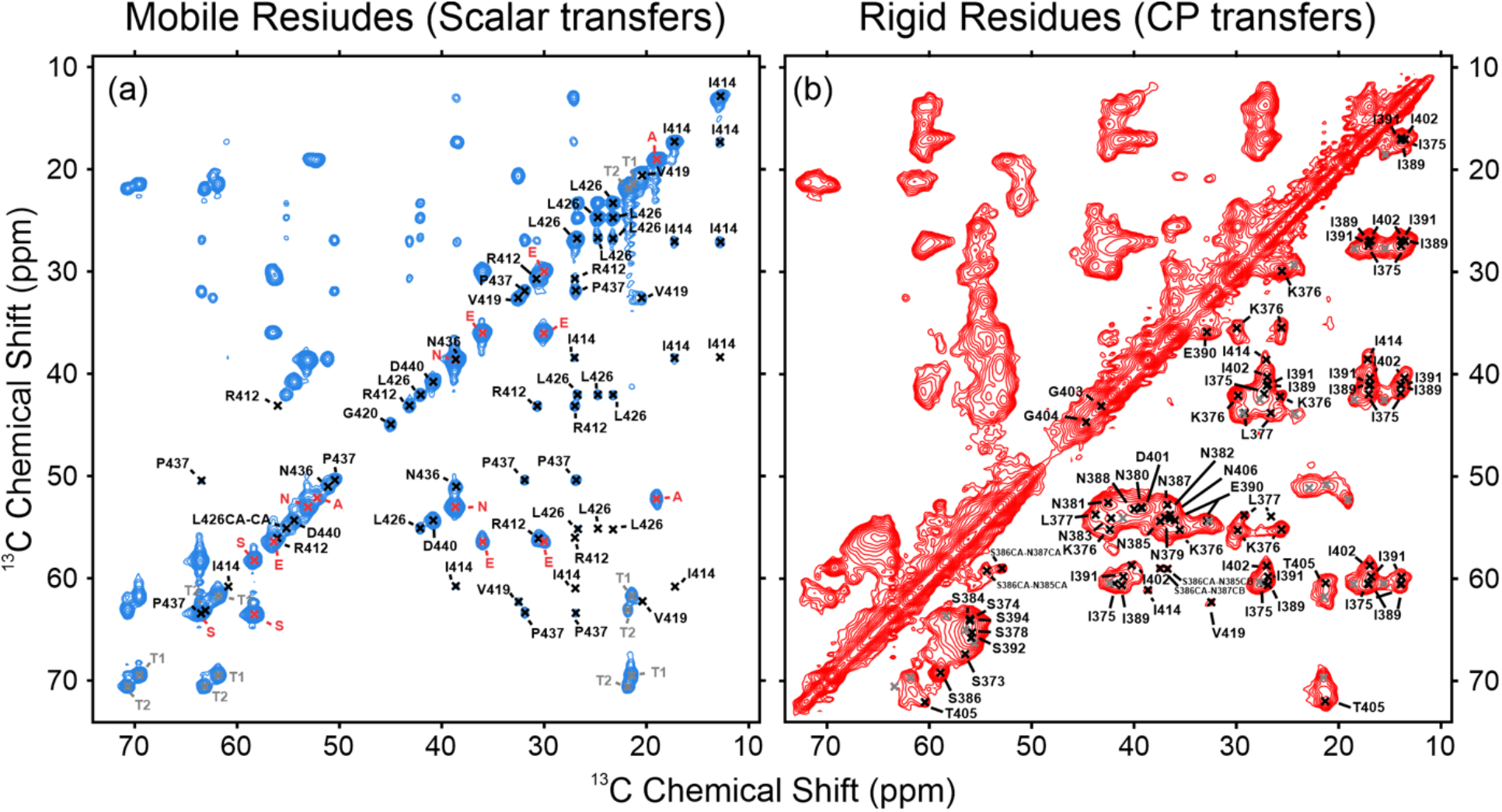
Solid State NMR of Tm1-LC Fibrils. (a) INEPT-TOBSY spectrum of the Tm1-LC fibrils reporting on residues exhibiting µs or smaller rotational correlation times. Resonances with unambiguous residue assignments are labeled in black, tentative resonance assignments for the ambiguous, degenerate residues in the C-terminal half of Tm1-LC are labeled in red, and the unassigned resonances are labeled in grey. (b) The aliphatic region of the cross polarization based ^13^C–^13^C DARR spectrum recorded with a 50 ms mixing time, reporting on relatively rigid residues in the fibrils. Resonances with unambiguous residue assignments are labeled in black and the unassigned resonances are labeled in grey.

**Figure 4.**
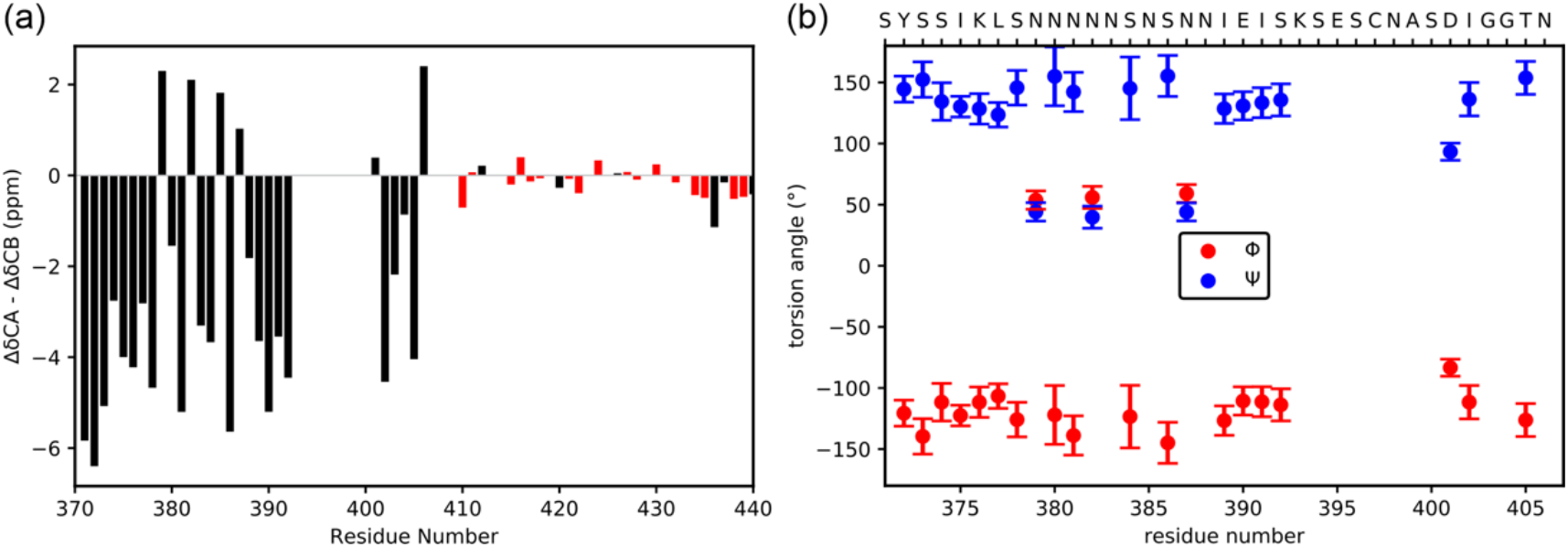
Secondary Structure Information Derived from the Solid State NMR Measurements. (a) Secondary chemical shifts relative to the random coil chemical shifts obtained from the Poulsen IDP calculator for the assigned residues from the cross polarization and scalar coupling based experiments. Red bars represent residues with tentative assignments from INEPT-TOBSY spectrum. (b) The φ and ψ torsion angle predictions from the TALOS-N analysis of the assigned chemical shifts from the cross polarization experiments. The error bars represent the standard deviations from TALOS-N.

Cross polarization based solid state NMR measurements report on the rigid residues in the Tm1-LC fibrils. Signals in these spectra arise from residues with ms or longer rotational correlation times representing the immobilized core of the fibrils. Unambiguous sequence specific assignment from measured chemical shifts of the rigid regions of the Tm1-LC fibrils are made with the assistance of the Monte Carlo simulated annealing algorithm MCASSIGN2b (Tycko, 2015) using data from 3D NCACX, NCOCX, and CANCO experiments. A complete description of the assignment process and the MCASSIGN2b parameters are described in the methods section. Sequence-specific assignments were obtained for residues SY-S373–K393 and D401–N406. K393 only has a ^15^N backbone assignment and no assigned ^13^C shifts. Supplementary Figure 6 shows 2D spectra from NCACX and NCOCX experiments with the resonance assignments labeled. Supplementary Figure 7 shows example 2D planes from the 3D experiments tracing out the connection between residues S373–I375. The resonance assignments are depicted on the 2D cross polarization based ^13^C–^13^C DARR spectrum in Figure 3b. For the signals observed in the “i” type NMR spectra (i.e. NCACX) reporting on rigid segments of Tm1-LC fibrils 72.5% have been assigned. Some of the unassigned peaks likely correspond to the residues between K393 and D401, although they could not be unambiguously placed on the sequence. This is due to either redundancy in ^15^N chemical shifts for multiple signals (nearly the same chemical shift value limiting the ability to match one signal to another), or a lack of a matching pair of CA and CB signals between the NCACX and NCOCX spectra. Additionally, one of the unassigned peaks likely corresponds to an alternative isoform for the sidechain, but not the backbone, of I375. This peak has nearly identical shifts for its N, CA, and CO atoms, although slight differences in sidechain ^13^C shifts. Two potential assignments for I375 in the solid state NMR data may correspond to different fibril quaternary structures observed in the cryo-EM data. Other than for I375, multiple assignments for all other rigid core residues were not consistent with the solid state NMR spectra. The lack of unique assignments for each fibril superstructural polymorph is likely due to two reasons: first, the cryo-EM class A–C models show excellent agreement up to the vicinity of residue K393, and second, solid state NMR linewidths observed here are likely not sufficiently narrow to tease out any differences in chemical shifts resulting from the nearly identical fibril structures.

Torsion angle predictions are generated from the assigned NMR chemical shifts for the rigid core of the Tm1-LC fibrils using the TALOS-N program are shown in Figure 4b (Bax, 2013). These torsion angles are then compared with the mean torsion angles across all five monomer conformations from the cryo-EM reconstructions. The torsion angle difference between the solid state NMR and cryo-EM analyses in Figure 5a demonstrate excellent agreement between the two methods. The uncertainties in Figure 5a are the TALOS-N prediction standard deviations. These uncertainties show that, for many rigid-core residues, the difference in torsion angle value between cryo-EM and solid state NMR predictions is close to or indistinguishable from 0°. In other words, the uncertainty is larger than the difference between the two methods. Numerically, across all the ψ and φ angles compared between cryo-EM and solid state NMR, the mean difference is −10°, while the average error from the TALOS-N prediction is ± 20°.

**Figure 5.**
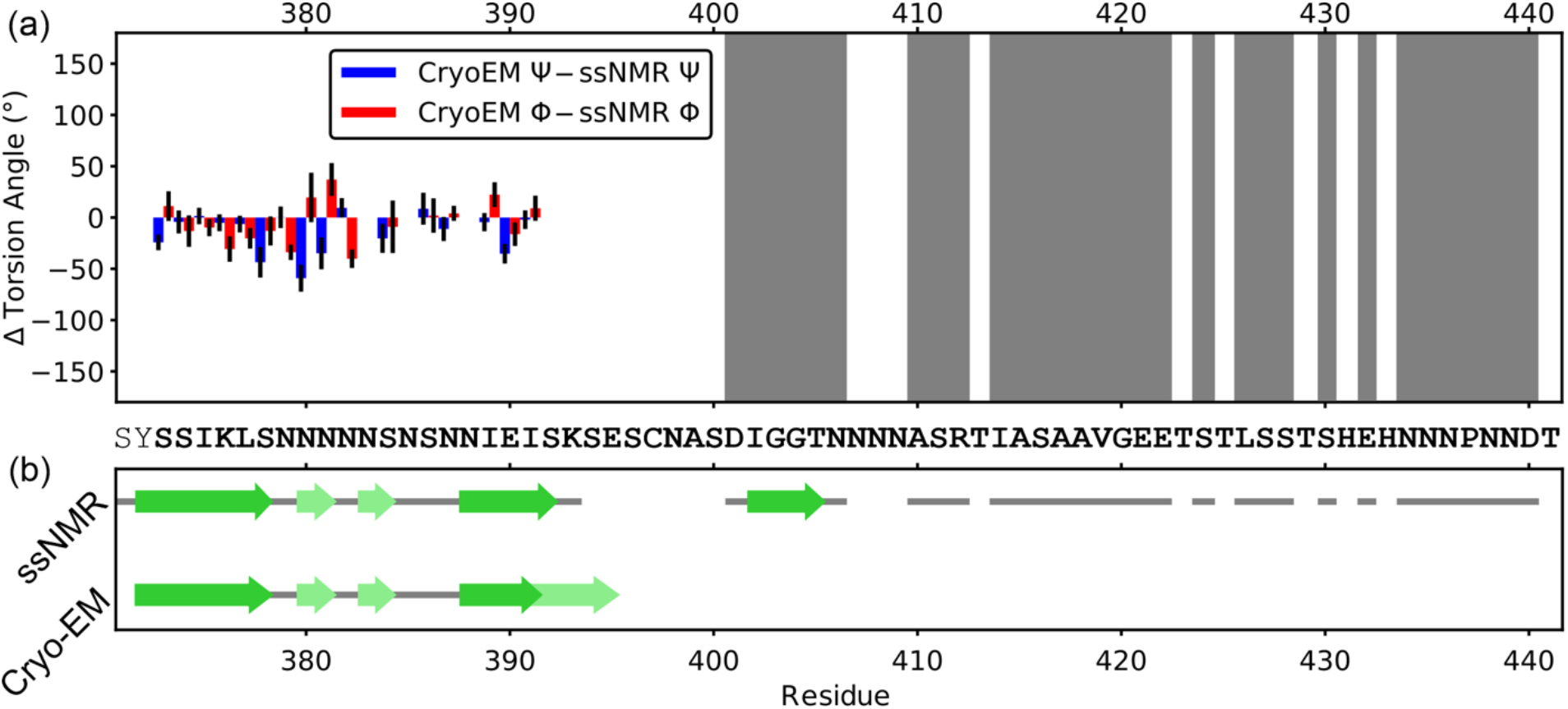
Comparison of Secondary Structure Between Solid State NMR and Cryo-EM. (a) Backbone torsion angle comparison represented as the difference between the mean angle across all five modeled Tm1-LC chains and the solid state NMR predictions from TALOS-N. The mean and standard deviation of the difference in torsion angles between cryo-EM and solid state NMR for all ψ and φ angles measured is −10° ± 20°. Each amino acid has a red bar for the φ angle and a blue bar for the ψ angle. The grey regions represent stretches of residues characterized exclusively by solid state NMR measurements. The black error bars are the standard deviations reported by the TALOS-N prediction. (b) Consensus secondary structure from the cryo-EM models compared with the secondary structure derived from the solid state NMR secondary chemical shifts and TALOS-N torsion angle predictions. The dark green arrows represent β-strand regions, light green arrows represent β-strand-like regions in the turn or β-strands not observed in all cryo-EM classes, and the solid gray lines represent observed amino acids without well-defined secondary structure, but includes both rigid and mobile regions of the Tm1-LC.

The torsion angles, along with the secondary chemical shifts for the NMR data, lead to the secondary structure cartoon models depicted in Figure 5b. Nearly the same regions are observed to be in a β-strand arrangement based on the cryo-EM and solid state NMR analysis. Both methods indicate a β-strand from residues S373–S378. The cryo-EM classes A–C show a nearly identical β-strand from residues N388–I391 in agreement with the TALOS-N solid state NMR analysis. However, all chains modeled in classes A–C, except one of the two chains in the class C repeating unit, the β-strand extends to E395. The TALOS-N prediction extends to residue S392. The lack of unambiguous assignments in the solid state NMR data from K393–S400 may be due to the structural heterogeneity in the Tm1-LC fibrils past K393, since the cryo-EM models shown in Figure 2c show structural divergence starting from this position. It is possible that some of the unassigned peaks in the solid state NMR data may stem from this structural heterogeneity. It should also be noted that for residues N379–N387 both solid state NMR and cryo-EM indicate a rigid turn region with some β-strand-like structure. In summary, the rigid core region consistently observed in all cryo-EM maps was also unambiguously assigned in the analysis of the cross polarization based solid state NMR experiments reporting on the rigid core of the fibrils.

In addition to the agreement between the cryo-EM and the solid state NMR determination of the rigid core residues, these NMR measurements additionally report on regions not observed in the cryo-EM reconstruction. Highly mobile residues in random coil conformations are identified for residues A410– T441 by INEPT, scaler coupling based measurements. Furthermore, the cross polarization based NMR measurements identify an additional rigid segment, D401–N406, in the fibrils containting a β-strand composed of residues I402–T405. The lack of this C-terminal β-strand in the cryo-EM density could be due to this additional β-strand interacting with the fibril core in multiple rigid arrangements, or this region may not strongly associate with the fibril core and remains only partially immobilized. Either situation would not result in well-defined density in the cryo-EM reconstruction.

An alternative explanation for the additional β-strand from solid state NMR is that these assignments come uniquely from a class D fibril conformation with a distinct rigid core compared to classes A–C. However, this contradicts the rigid body fitting of the class B model into the class D density map in supplementary figure 4, although the low resolution for class D precludes ruling out the participation of other residues. A previous solid state NMR study on His-tagged Tm1-LC fibrils indicated β-strand secondary structure for residues N398–S411, which overlaps partially with the solid state NMR analysis in this work indicating β-strand secondary structure for I402–T405 (Sysoev, 2020). However, the His-tagged Tm1-LC fibrils have different NMR chemical shifts for residues I402–T405 than the those determined here, suggesting this region adopts distinct structural conformations depending on the nature of the N-terminal region of the Tm1-LC. Therefore, our work in the context of previous Tm1-LC fibril structural data shows that Tm1-LC has potentially multiple fibril-forming core regions, whose rigidification is construct-dependent. Overall, solid state NMR data add to the structural characterization of the rigid regions of Tm1-LC fibrils, and provides a characterization for residues that are unable to be modeled from cryo-EM data.

## Discussion

Cryo-EM and solid state NMR structural characterization of SY-tagged Tm1-LC fibrils show excellent agreement demonstrating the robustness of both techniques to determine the fold of the rigid fibril core region. TALOS-N torsion angle predictions and secondary chemical shifts from solid state NMR agree with the structural models from cryo-EM. In addition, the solid state NMR analysis identifies an additional β-strand region, and also determines that the C-terminus of the protein is highly mobile and in a random coil conformation.

Compared with solid state NMR, cryo-EM possesses some distinct strengths to study protein fibril structures. Without additional samples and experimentation, solid state NMR was unable to determine that the samples contained superstructural polymorphs of Tm1-LC, while cryo-EM shows the presence of four unique 2D classes with classes A–C having nearly identical tertiary structures but different quaternary structures. The lack of unique chemical shifts for each molecule and class of Tm1-LC fibrils in the solid state NMR data, despite different packing interface regions in each monomer, is unsurprising given the nearly identical torsion angles of each molecule in the repeating fibril unit (Figure 2d). Torsion angles directly influence the chemical environment around the atoms in each residue, which largely determines the NMR chemical shifts. Therefore, the chemical shifts arising from each protein molecule in the fibril structure are expected to be indistinguishable based on the similar tertiary structure observed in the cryo-EM map, at least given the NMR experiments used in this work (de Dios et al., 1993; Spera et al., 1991). Additional solid state NMR experiments measuring intermolecular interactions could permit the identification of superstructural polymorphs and determine fibril registry, as has been demonstrated for other fibrils (Murray, 2017; Tycko, 2007).

Despite not observing the quaternary structure variations observed using cryo-EM, a limited solid state NMR dataset obtained on a single uniformly ^13^C and ^15^N labeled sample like the one used here has its own advantages for the structural analysis of protein fibrils. First, different degrees of motion are readily characterized, since the rigid fibril core observed here is shown to have only small amplitude motions on the time scale of ms or slower and the flanking regions are shown to have rapid motion on the time scale of µs or faster. Second, solid state NMR can detect structured regions that are invisible in the cryo-EM map, as our analysis demonstrates β-strand character for I402–T405, which was not observed in any of the cryo-EM based-models. Third, the secondary structure of mobile protein regions is characterized more completely with solid state NMR, as we show here that the residues outside the fibril core are not adopting structurally heterogeneous rigid conformations, but are undergoing fast reorientations and lack well-defined secondary structure. Cryo-EM efforts have made progress in the detection of more mobile regions of proteins in the low-resolution regions of density maps, however, NMR analysis provides a direct residue-specific probe of highly mobile regions, especially for loops or intrinsically disordered domains that lack any significant density in cryo-EM reconstructions (Zhong et al., 2021). Finally, although this was not an issue for these Tm1-LC fibrils, solid state NMR does not require well-dispersed particles nor a helical twist to achieve structural detail at the level of individual amino acids and is therefore applicable to a range of systems not directly accessible by cryo-EM.

In the published literature, many cryo-EM but not solid state NMR studies require additional sample treatments such as proteases to cleave off intrinsically disordered regions or the addition of detergents to avoid particle clumping (Arseni, 2023, Arseni, 2021; Cao et al., 2019; Jiang et al., 2022; Schweighauser et al., 2022). If there is a biological importance to the flanking regions of the fibril core, a hybrid strategy would be of great benefit. For example, protease-treated fibrils studied with cryo-EM could identify the structural motifs leading to the formation of the fibril core conformation or conformations, and solid state NMR could then be used to characterize intact full-length fibrils untreated with proteases to gain an understanding of the structure and molecular motion of regions outside the rigid fibril core. Such a combined approach could lead to a more complete understanding of the biological question at hand by characterizing both the core and flanking regions of protein fibril assemblies.

The importance of flanking fibril core regions has been demonstrated in the literature. For example, the C-terminal region of the fused in sarcoma (FUS) N-terminal low complexity domain fibrils has been shown to conformationally select for a fibril conformation by acting as an entropic tail, thus stressing the importance of folded and unfolded regions in determining protein fibril structure (Kumar et al., 2021). Similarly, conformations adopted by SY-tagged Tm1-LC analyzed here are distinct from the conformations adopted by His-tagged Tm1-LC, despite evidence that the His-tag was not a rigid component in the fibril core (Sysoev, 2020). In addition to influencing the conformations adopted by the rigid regions of protein fibrils, non-rigid regions are important to characterize due to their functional role. For example, regions flanking the fibril core of the Orb2A protein have been shown to be important for its functional interaction with partner proteins (Soria et al., 2022). In this case, solid state NMR allowed direct characterization of the fuzzy coat region Orb2A fibrils (Soria et al., 2022).

This work has focused on the complementary nature of cryo-EM and solid state NMR studies of protein fibrils. Solid state NMR is able to study highly mobile and immobilized core regions for protein fibrils at the same time, whereas cryo-EM is highly robust at determining relatively static structures. It should be noted that more broadly, viral capsids, membrane bound proteins, and other large macromolecular complexes displaying a range of molecular motions across their structures or requiring variable cryo-EM grid freezing conditions and treatments for well-dispersed assemblies, can also be more thoroughly and readily characterized with a combination of cryo-EM and solid state NMR.

## Methods

### Purification of SY-Tm1C Protein

An expression plasmid for GFP-GDEVD-SY-Tm1-LC was constructed in pHis-parellel1-GFP vector as described previously (Kato et al., 2017; Sheffield et al., 1999). Tm1-LC contains residues 373–441 of the *Drosophila* Tm1-I/C isoform. GDEVD is a caspase-3 cleavage sequence. Tm1-LC does not have any tyrosine or tryptophan residues. Thus, the dipeptide SY is added in front of Tm1-LC so that the protein has measurable absorbance at 280 nm for protein quantification. PCR DNA fragments for GDEVD-SY-Tm1-LC were amplified from the full-length Tm1-I/C isoform as a template and sub-cloned into the pHis-parallel1-GFP vector. The Tm1-LC sequence was confirmed by standard commercial DNA sequencing methods.

The plasmid was transformed in *E. coli* BL21(DE3) cells. Expression and purification of the protein is carried out as described previously (Kato, 2017). Briefly, the fusion protein was overexpressed by induction with 0.5 mM isopropyl β-D-1-thiogalactopyranoside (IPTG) at 20 °C overnight. Cells were harvested and resuspended in lysis buffer containing 50 mM tris(hydroxymethyl)aminomethane (Tris-HCl) pH 7.5, 500 mM sodium chloride, 10 mM β-mercaptoethanol (BME), 2 M urea and one tablet of EDTA-free protease inhibitor cocktail (Sigma). Cell suspensions were lysed by sonication for 2 minutes (10 seconds on and 30 seconds off) on ice. Cell lysates were centrifuged at 142,000 x g for 1 hour at 4 °C. Supernatants were mixed with Ni-NTA resin (Gold Bio), shaken gently for 15 min in a 4 °C cold room, then poured into a glass column. The resin was washed with buffer containing 20 mM Tris-HCl pH 7.5, 500 mM sodium chloride, 10 mM BME, 0.1 mM phenylmethylsulfonyl fluoride (PMSF), 2 M urea and 20 mM imidazole. Bound proteins were eluted from the resin with an elution buffer containing 20 mM Tris-HCl pH 7.5, 500 mM sodium chloride, 10 mM BME, 2 M urea and 250 mM imidazole. Purified proteins were concentrated using Amicon Ultra filters (Millipore) to a concentration of 60 mg/ml.

Tag-free SY-Tm1-LC was produced by cleavage of the GFP tag with caspase-3 protease. Briefly, the concentrated protein was diluted to 2–3 mg/ml in a gelation buffer containing 20 mM Tris-HCl pH 7.5, 200 mM sodium chloride, 0.5 mM ethylenediaminetetraacetic acid (EDTA), 0.1 mM PMSF and 20 mM BME. Caspase-3 protease was then added at a 1:2000 ratio to the target protein. The reaction mixture was gently rotated at room temperature (RT) overnight. Cleaved proteins were mostly precipitated during the cleavage reaction and recovered by centrifugation at 4000 rpm in a tabletop microfuge for 30 min at 4 °C. The precipitated proteins were dissolved in 2 ml of a denaturing buffer containing 20 mM Tris-HCl pH 7.5, 200 mM sodium chloride, 10 mM BME, 0.5 mM EDTA, 0.1 mM PMSF and 6 M guanidinium-HCl (GdnHCl), and then loaded onto a Superdex S-200 gel filtration column equilibrated with 20 mM Tris-HCl pH 7.5, 200 mM sodium chloride, 10 mM BME, 0.5 mM EDTA and 2 M GdnHCl. Fractions containing the cleaved proteins were combined and concentrated to ∼60 mg/ml. Powder of GdnHCl was added to the protein solution to make the final GdnHCl concentration of 6 M, and samples were stored at −80 °C.

### Fibril Formation

For uniformly labeled ^13^C ^15^N tag-free SY-Tm1-LC fibrils for NMR analysis, the protein in 6 M GdnHCl stock solution was diluted in 300 ml of the denaturing buffer to a final protein concentration of 5 mg/ml and then dialyzed against the gelation buffer overnight. The protein solution was recovered in microcentrifuge tubes. 10% sodium azide solution was added to a final concentration of 0.02%. The protein solution was incubated at 4 °C for 3 d, then a brief sonication was applied to facilitate polymerization. It was incubated at 4 °C for 4–5 more days and checked by negative-stained TEM for fibril formation. Non-isotopically labeled protein for cryo-EM analysis was generated identically, except at smaller scale. In brief, natural abundance tag free SY-Tm1-LC in 6 M GdnHCl stock solution was diluted in 1 mL of denaturing buffer to a final protein concentration of 5 mg/mL and then, the fibril preparation procedure was identical to isotopically labeled fibrils.

### Negative Stain TEM Imaging

Ten µl of fiber solution was applied to carbon film grids (CF200-Cu, Electron Microscopy Sciences) for 1 min and removed by blotting with a laboratory tissue. Grids were briefly washed with 5 mM EDTA pH 8.0, exposed to 2% uranyl acetate solution for 15 s and air-dried. TEM images were obtained with Tecnai Spirit electron microscopes at 120 kV.

### Cryo-EM Grid Preparation and Data Collection

The Tm1-LC fibril sample was prepared at 1 mg/ml concentration by dilution of fibrils with gelation buffer. Three µl of the sample was applied to Quantifoil 300 mesh R1.2/1.3 grids (Quantifoil, EMS) that were glow discharged using a Pelco EasiGlo. Grids were flash-frozen in liquid ethane using an FEI Vitrobot mark IV (Thermo-Fisher Scientific). The settings of the Vitrobot were blot force −5, blot time 3 s, relative humidity 95%, and 4 °C. Sample grids were screened on a Talos Arctica microscope before data collection. The cryo-EM data were acquired on a Titan Krios microscope at the UT Southwestern Cryo-Electron Microscopy Facility, equipped with BioQuantum energy filter and K3 direct electron detector. 15,219 movies were acquired in TIFF format at a pixel size of 0.415 Å in super-resolution mode using SerialEM (Mastronarde, 2005). The accumulated total dose was 52 e^-^/Å ^2^ which was fractionated into 50 frames. The defocus range of the images was set to be −1.0 to −2.4 μm.

### Helical Reconstruction

Movie frames were motion-corrected and averaged into single images using the MotionCor2 program (Zheng et al., 2017). The CTF parameters were calculated using Gctf (Zhang, 2016). Manually picked fibrils from 15 images were used for neural network training in crYOLO (Wagner et al., 2019). The trained model was then used to auto-pick the whole dataset in crYOLO and the coordinates were imported into RELION. All the rest of the data processing steps were carried on in RELION. 2,295,590 filament segments were extracted at 768 box size, with an inter-box distance of 14.25 Å, and were used for one round of 2D classification. Four distinct types of fibrils were identified based on the features of the 2D classes, containing 1,401,970 segments in total. For each type, 2D classes with obvious cross-over features were selected and used for initial model generation using relion_helix_inimodel2d, with the cross-over distance estimated from these 2D classes (Scheres, 2020). The segments were then reextracted at 320 box size, followed by additional 2D classifications, 3D classifications, and 3D refinements. The number of segments used for the final reconstruction, resolution reported by RELION post-processing, and the refined helical parameters are listed in Table 1. The cryo-EM density map for class A, which has the highest resolution, has been deposited in the EMDB (EMD-45130), to be released upon publication.

### Model Building and Refinement

The SY-Tm1-LC class A model was first manually built using the Coot program, referring to the CA trace model from the program Model Angelo (Emsley, 2010). Then, the other class models were manually built, referring to the class A model. The models were then refined with the real-space refinement mode of the Phenix program (Afonine et al., 2018). The statistics of the model validations are summarized in Table 2 and Supplementary Tables 1–2. The final structure of the best class, class A, is deposited in the PDB (9C1U), to be released upon publication.

### Torsion Angle Calculations from the Cryo-EM Structure

Torsion angles from the Tm1-LC monomers fit into the cryo-EM density map were independently measured by loading the structure into ChimeraX and using the ‘torsion’ command for each residue to yield the values plotted in Figure 2d, Supplementary Figure 5, and Figure 5a (Pettersen et al., 2021). RMSD values are calculated in ChimeraX with the use of the “align” command to first optimize the alignment of the two molecules and then calculate RMSD (Pettersen, 2021). The atoms used for the alignment are stated in the main text and Supplementary Tables 3–5. Rigid body fitting of class B into one half of the cryo-EM density map from class D was calculated with the “fitmap” command, specifying only backbone atoms CA, C, O, and N to be used in the fitting.

### Solid State NMR

The assigned observed ^13^C and ^15^N NMR chemical shifts have been deposited in the Biological Magnetic Resonance Bank (BMRB 52468). Tm1-LC fibrils were iteratively packed into a thin-walled 3.2 mm pencil style zirconia rotor (Revolution NMR) using a metal spatula and ∼5 s centrifugation steps at < 3,000 g. The rotor was then centrifuged at 20,000 g for 1.75 hr at 8 °C using a home-built device to hold the rotor in the centrifuge tube. The rotor drive tip and top cap were attached using cyanoacrylate glue gel to seal the sample in the rotor and maintain hydration.

An 18.8 T Bruker Avance III NMR spectrometer was used for all experiments. The spectrometer was operated with a triple resonance BlackFox probe equipped with a ^1^H loop gap resonator to eliminate sample heating due to high power ^1^H decoupling. The magic angle spinning rate (MAS) was 12.75 kHz, the cooling air set to 243 K, and heater set to 263 K. The combination of cooling air, heater settings, and MAS rate provided a sample temperature of ∼10 °C based on calibration of the system with ^79^Br (Thurber et al., 2009). Indirect data acquisition was recorded using States-TPPI mode for the cross polarization-based experiments. The indirect data acquisition for the INEPT-TOBSY experiment was recorded in TPPI mode. The spectra were referenced to the DSS scale using an external 1-^13^C labeled alanine sample set to 179.8 ppm. A detailed list of all parameters used in the NMR experiments is found in Supplementary Table 6.

### The Timescale of Motion for the TOBSY Signals

The slowest time scale of motion observable in the INEPT-TOBSY experiment was calculated by solving the following formula for the correlation time, τ_c_, using a T_2_ value equal to the 1.0 ms J-delay used during the INEPT transfer periods. The formula assumes the ^1^H-^13^C dipolar coupling is the main component contributing to transverse relaxation under 12.75 kHz MAS (Bloembergen et al., 1948).

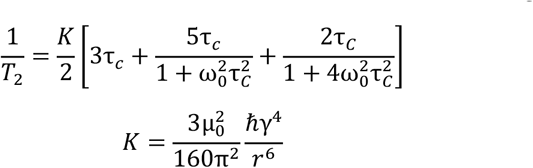

where T_2_ = 1 ms, ω_0,13C_ = 201.197878 MHz, µ_0_ = 1.2566·10^−6^ N/(Amp^2^), γ_13C_ = 67.2828·10^6^ rad/(s·T), and r = 1.09 Å (the typical H-C bond length). The calculated correlation time, τ_c_, is 1.6 µs. Therefore, for a region of the protein with a τ_c_ 1 µs, ∼37% of the signal remains after each half-echo in the INEPT transfer period. Regions of the protein with correlation times faster than 1 µs would therefore retain observable signal intensity.

### Solid State NMR Assignments

Resonance assignments are made with a combination of the information from all the spectra. The rigid sites are assigned with the Monte-Carlo annealing algorithm MCASSIGN2b (Tycko, 2015), using the chemical shift peaks from the 2D/3D NCACX and NCOCX spectra, and the 3D CANCO spectra. Input chemical shift tables are in the Supplementary Tables 11–13. The MCASSIGN2b program is ran with parameters for “good”, “bad”, “edge”, and “used” of, respectively, (0, 10), (10, 60), (0, 8), and (0, 2) over 200 steps with 10^7^ iterations per step, and 50 independent runs with unique random number seeds.

The calculation resulted in unambiguous assignments for residues K376, S378–I391, and D401– N406. Large uncertainties were used in the input chemical shift tables to yield only the most certain assignments first, therefore, out of the 39 signals initially in the NCACX MCASSIGN2b table, 20 remaining signals were not yet unambiguously assigned. Additional assignments were made using a manual assignment procedure analogous to the solution NMR method to allow for unambiguouos assignments for residues, in sum with the MCASSIGN2b assigned residues, for: SY-S373–K393, and D401–N406. All unambiguous assignments from CP-based experiments are listed in Supplementary Table 7. The remaining 11 unassigned peaks are listed in Supplementary Table 9. An example of how the signals are linked together for the manual assignments from the 3D spectra is depicted in Supplementary Figure 7.

To determine if the unassigned NMR chemical shifts were consistent with an alternate conformation for the Tm1-LC in the sample, the remaining signals were used as input for the same manual assignment procedure. The entire Tm1-LC sequence was considered in the procedure. None of the unassigned signals were assigned to residues SY-S373–I393 or D401–N406 or anywhere else in the Tm1-LC amino acid sequence. Apart from I375 having potentially an alternative sidechain conformation, we do not observe two sets of signals for each of the two distinct monomers in the Tm1-LC fibril structure. One set of signals is observed in the spectra recorded for this study despite the superstructural polymorphs identified by cryo-EM.

Assignments for the dynamic portions of the Tm1-LC fibrils were made as follows. For many residues in the TOBSY spectra in Figure 3a, all aliphatic ^13^C chemical NMR shifts were measured, allowing unambiguous assignment of amino acid residue type based on random coil chemical shifts. These amino acid types corresponded to the unique residue amino acids in the unassigned, C-terminal, region of the Tm1-LC amino acid sequence, providing unambiguous assignments for R412, I414, V419, G420, L426, N436, P437, and D440. These assignments all had chemical shifts consistent with an IDP and were scattered throughout the C-terminal half of the sequence, see Figure 3a. Therefore, the Poulsen IDP calculator, which includes nearest neighbor effects, was used to estimate the chemical shifts for the remaining residues in the region A410–T441(Kjaergaard, 2011). The estimated CA and CB NMR chemical shifts for the Ala, Ser, Glu, and Asn residues in the C-terminal end of the Tm1-LC sequence are indistinguishable given the linewidths in the TOBSY spectra. Therefore, the presence of a single set of ^13^C resonances in the TOBSY spectra for multiple A, S, E, and N residues yielded the tentative assignments that are represented in red in the spectrum in Figure 3a and the secondary chemical shift plot in Figure 4a. Supplementary Table 8 contains the assignments from the TOBSY-based spectra, and Supplementary Table 10 has the unassigned signals from the TOBSY-based spectra.

### Secondary Chemical Shifts and Torsion Angles

The Poulsen IDP calculator was used with the complete SY-S373–T441 Tm1-LC amino acid sequence to determine the expected random coil shifts (Kjaergaard, 2011; Kjaergaard, 2011; Maltsev, 2012; Schwarzinger, 2001). The difference between observed and calculated random coil NMR chemical shifts yielded secondary chemical shifts, Δδ_CA_ and Δδ_CB_. The difference between these two differences is plotted in Figure 5a.

TALOS-N was ran with standard parameters and the assigned chemical shifts to generate predictions for the φ and ψ torsion angles (Bax, 2013). “Strong” and “Generous” predictions are included in the plot in Figure 4 and Figure 5, and the TALOS-N output standard deviations are used as errors. TALOS-N predicted secondary structure motifs corresponds to β-strand for residues S373–L378, I389– S392, coil/turn for the SY tag, N379–N388, and D401–N406, and no presence of helical structure. Note that for this sample, purely using the secondary structure predictions (the TALOS-N “predSS” output) is a more conservative estimate of secondary structure compared to the use of experimentally measured secondary chemical shifts. Therefore, using a combination of the information gained from torsion angles, secondary chemical shifts, and secondary structure predictions output by TALOS-N, we have determined the β-strand regions depicted in Figure 5b, which, in addition to those listed above directly from the TALOS-N secondary structure prediction, also includes I402–T405 as forming a β-strand motif. The lack of prediction by TALOS-N for this region likely is due to the G403–G404 motif having ill-defined torsion angle predictions, perhaps due to this motif being under sampled in the TALOS-N database of structures, despite the secondary chemical shifts for these residues clearly indicating β-strand character, with the large negative secondary chemical shifts seen in Figure 4a.

## Acknowledgements

We thank Dr. Ping Yu for help with the NMR spectrometers. The UC Davis NMR Core Facility was supported though awards NSF DBI-0722538 and NIH 1S10RR013871-01A1. DTM acknowledges support from NIGMS award 5R35GM142892. UC Davis also provided generous start-up funding for DTM. We thank the Structural Biology Laboratory and the Cryo-Electron Microscopy Facility at UT Southwestern Medical Center which are partially supported by grant RP220582 from the Cancer Prevention & Research Institute of Texas (CPRIT). The content is solely the responsibility of the authors and does not necessarily represent the official views of the National Institutes of Health.

