## Supplemental Information for "Cryo-EM and Solid State NMR Together Provide a More Comprehensive Structural Investigation of Protein Fibrils"

### Supplementary Information

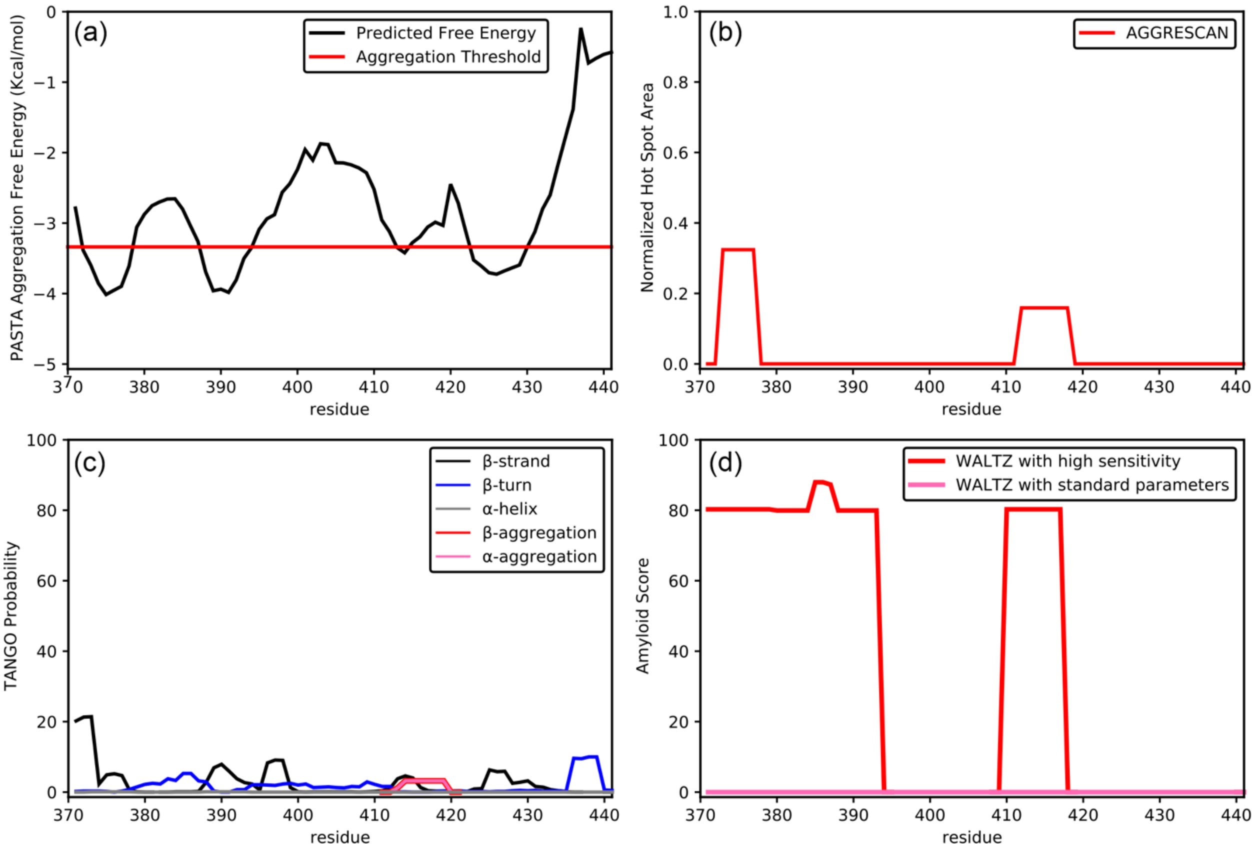

**Supplementary Figure 1. Aggregation Prediction Programs do not Strongly Predict the Rigid Core of the SY tagged Tm1-LC Fibrils.** (a) Free energy of aggregation predicted by PASTA 2.0 (Walsh et al., 2014). The calculation was run with standard parameters, and the 90% specificity option. The only regions predicted to aggregate are short in sequence, therefore no high-probability aggregation is predicted, see Figure 1 in the main text. (b) The AGGRESCAN prediction (Conchillo-Solé et al., 2007), run with standard parameters, has only weakly predicted aggregation regions. The first region, S373–L377, partially overlaps with the fibril core observed by solid state NMR and cryo-EM. (c) TANGO prediction (Fernandez-Escamilla et al., 2004; Linding et al., 2004; Rousseau et al., 2006) of secondary structure and aggregation displays a lack of both. (d) WALTZ prediction (Maurer-Stroh et al., 2010) using the standard parameters does not predict any amyloid formation for the Tm1-LC. Using WALTZ's less accurate, more sensitive parameters, Tm1-LC is predicted to form an amyloid; the first region is somewhat consistent with the core experimentally observed using solid state NMR and cryo-EM.

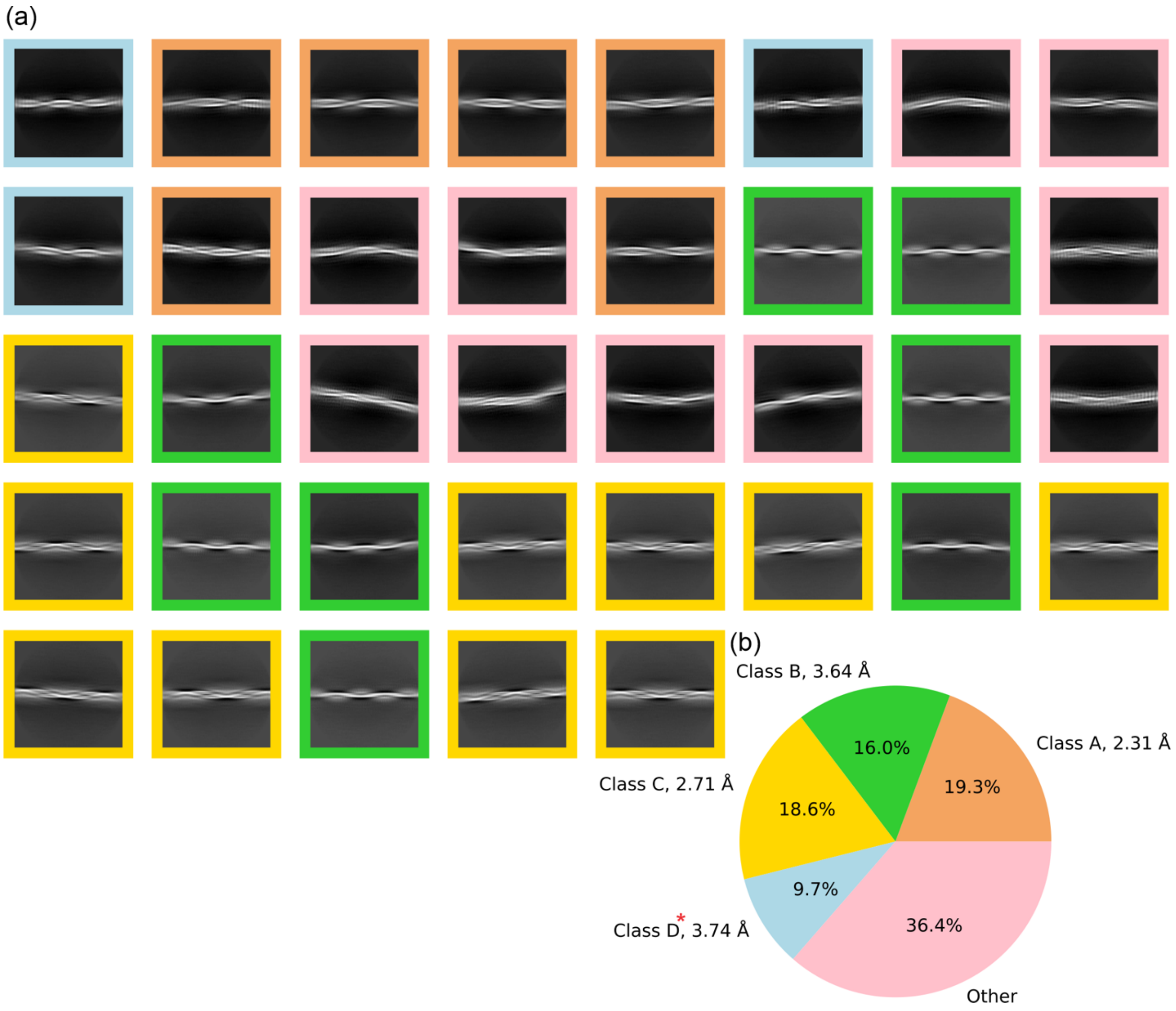

**Supplementary Figure 2. Cryo-EM 2D Classification.** (a) 2D classification in RELION shows four distinct classes. The border color corresponds to (b), reproduced from Figure 2. (b) Particle count percentages from the overall data collection. "Other" represents particles without consensus features. The red asterisk denotes the overestimate in resolution from class D resulting from the helical spacing observed in the 3D refined map.

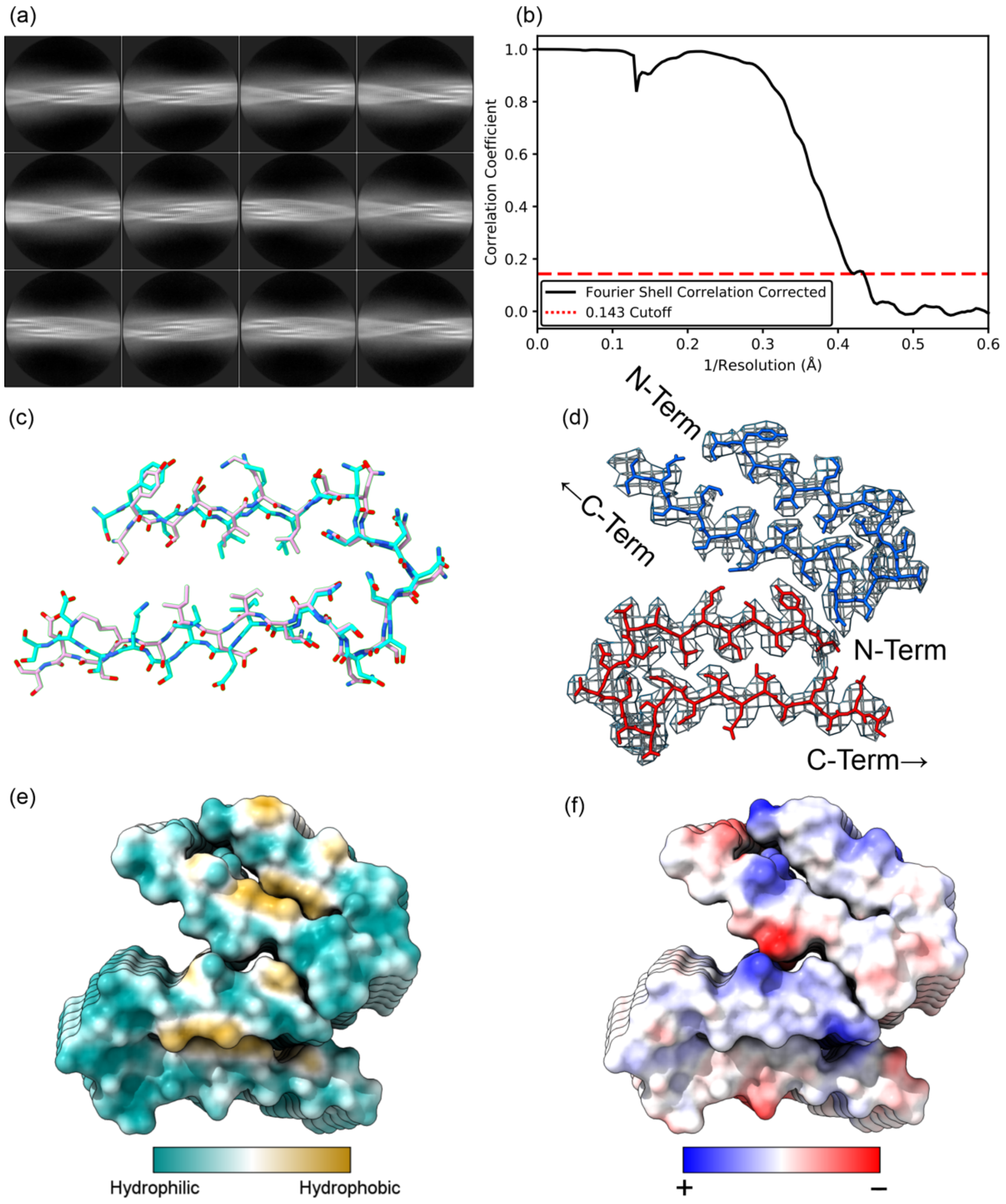

**Supplementary Figure 3. Class A Fibril Structure Analysis.** (a) Representative 2D classes from the cryo-EM reconstruction demonstrate the fibrils have a clear twist and a cross-β morphology. (b) FSC curve output from RELION. (c) The aligned overlay between the two unique monomer folds in the repeating fibril unit shows a near-identical structure. (d) Class A cryo-EM density with the modeled chain. Density is set to level 0.00807 with a ChimeraX step size of 2 (Pettersen et al., 2021). (e) Hydrophobicity plot of the class A model calculated with ChimeraX (Pettersen, 2021). (f) Electrostatic plot calculated from the class A model with ChimeraX (Pettersen, 2021).

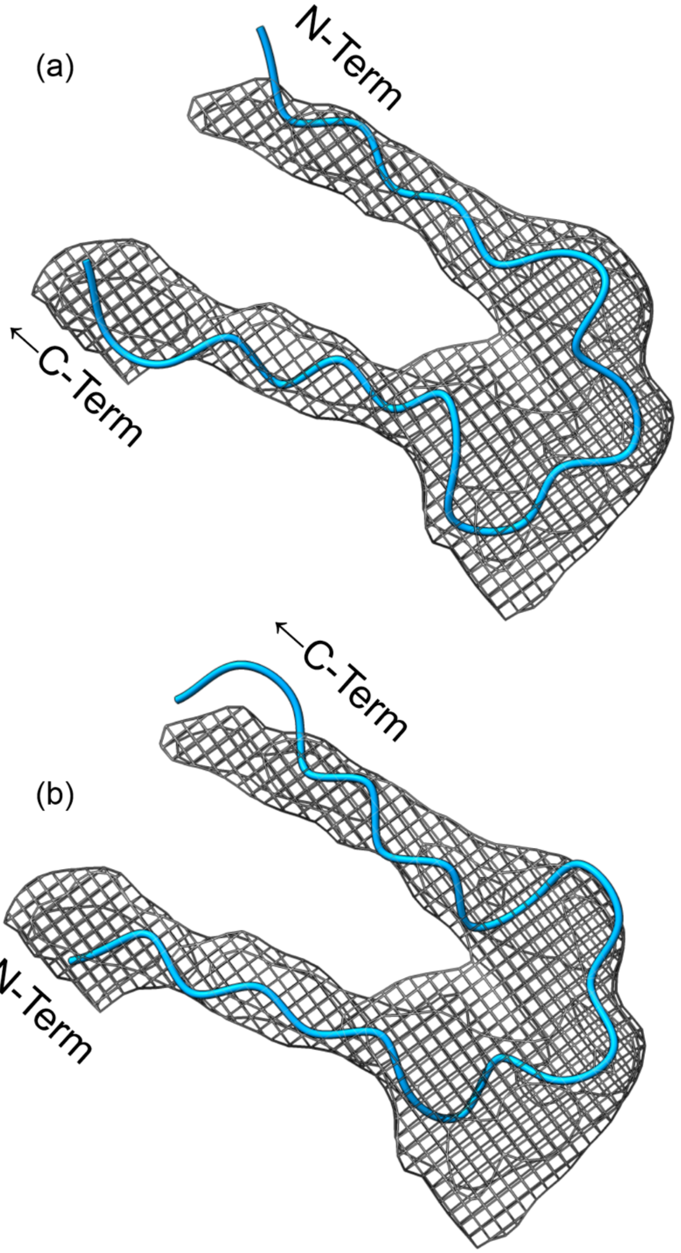

**Supplementary Figure 4. Class D shows a Similar Structure.** (a) and (b) Class B rigid body fitting into the Class D cryo-EM Density Map, with the molecule rotated 180° in (b) relative to (a). This change in starting coordinates leads to essentially the same fit quality. ChimeraX (Pettersen, 2021) was used with the "fitmap" command with only backbone atoms, CA, C, O, and N, to find the overlap between the map and model. (a) has an average map fit value of 0.01344 and (b) 0.01226.

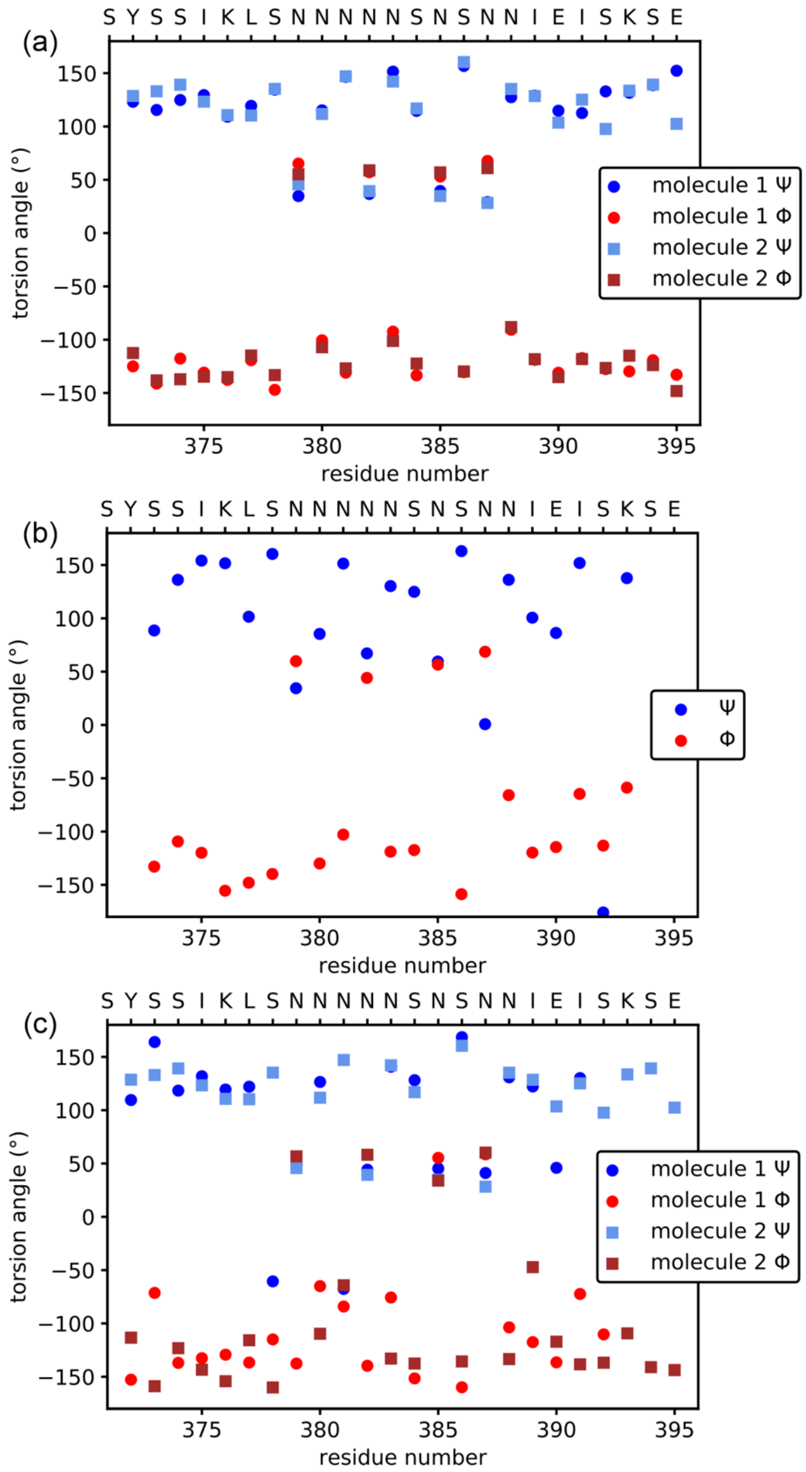

**Supplementary Figure 5. Torsion angles for Classes A–C.** Torsion angles from the cryo-EM modeled Tm1-LC protein chains. (a) Class A torsion angles, where molecules 1 and 2 represent the separate protein monomers in the helical repeating unit. (b) Class B torsion angles. (d) Class C torsion angles, where molecules 1 and 2 represent the separate protein monomers in the helical repeating unit. In the main text, Figure 2d is the combined overlay of all of these data.

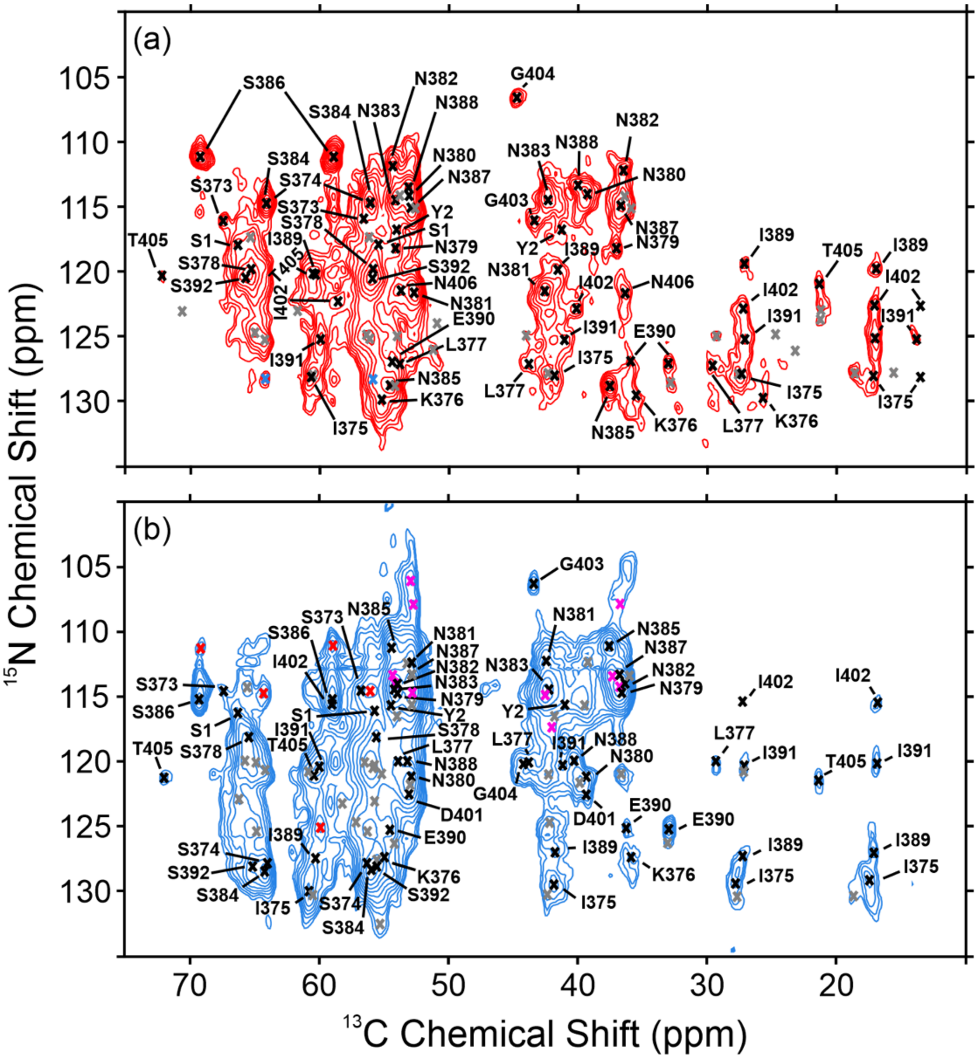

**Supplementary Figure 6. Solid State NMR Assignments Overlayed on 2D NCACX and NCOCX Spectra.** (a) Cross polarization based-NCACX spectrum with 50 ms DARR mixing, which gives rise to intraresidue cross peaks. Grey labels represent unassigned peaks, and blue labels represent interresidue, "i−1" cross peaks, i.e. the cross peaks appear with greater intensity in the NCOCX spectrum. (b) Cross polarization based-NCOCX spectrum with 50 ms DARR mixing, which gives rise to interresidue cross peaks between $^{13}C$ chemical shifts from the "i" residue and the $^{15}N$ chemical shift from the "i+1" residue. Labels represent the assignments from the $^{13}C$ shifts. Grey labels are unassigned, pink labels likely arise from transfers from the amide side chain of either Q or N residues, and red labels are from intraresidue cross peaks, i.e. they appear with greater intensity in the NCACX spectrum. Contours are drawn at intensity levels increasing by a factor of 1.3 for both spectra.

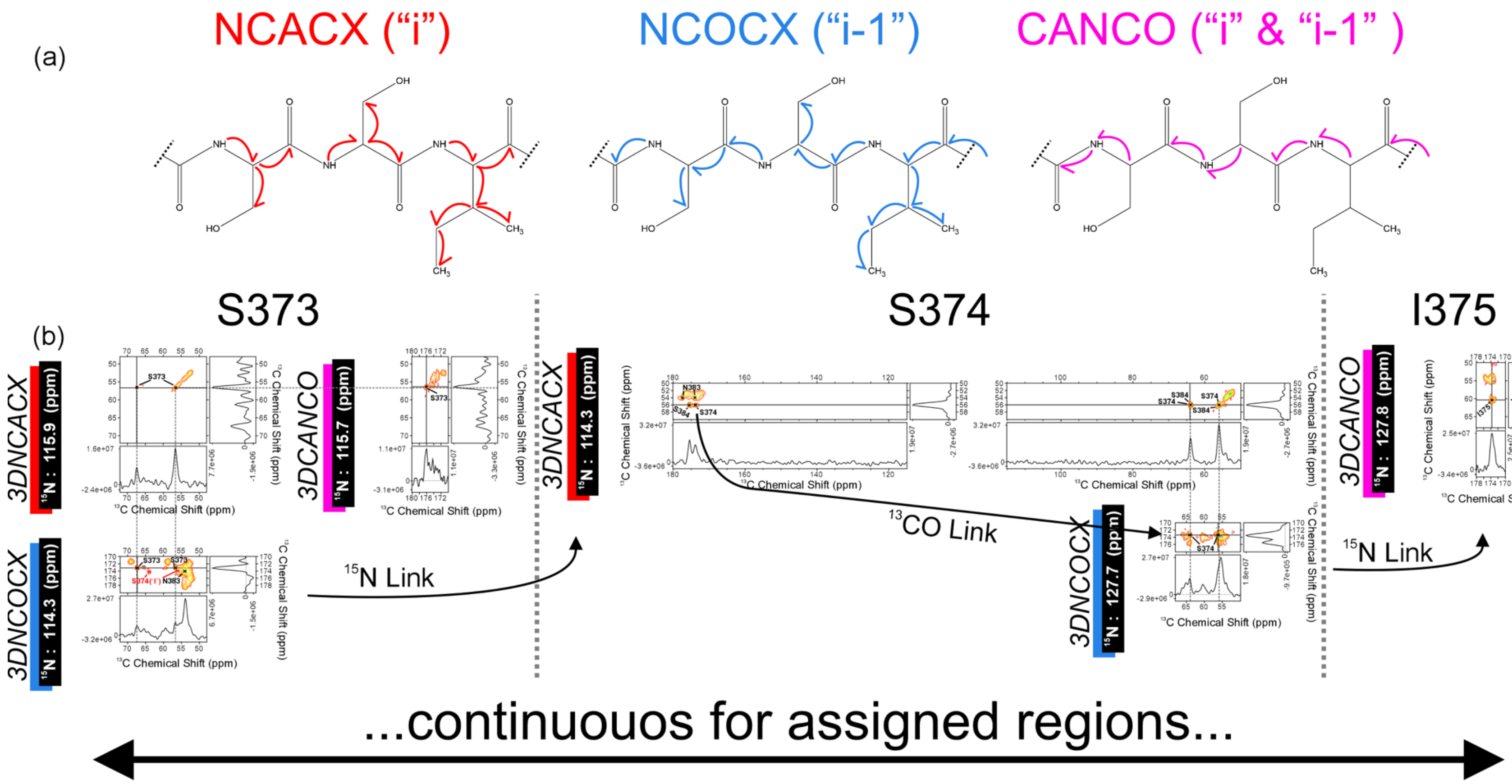
(a)
NCACX (“i”)
NCOCX (“i-1”)
CANCO (“i” & “i-1” )
(b)
S373
S374
I375
3DNCACX
15N : 115.9 (ppm)
3DNCOCX
15N : 114.3 (ppm)
3DCANCO
15N : 115.7 (ppm)
3DNCACX
15N : 114.3 (ppm)
3DNCOCX
15N : 127.7 (ppm)
3DCANCO
15N : 127.8 (ppm)
15N Link
13CO Link
15N Link
13C Chemical Shift (ppm)
...continuouos for assigned regions...

**Supplementary Figure 7.** (a) Diagram of the magnetization transfers between observed nuclei in the three cross polarization-based solid state NMR experiments used to make the assignments for the rigid residues within the fibrils. (b) Demonstration of the assignment process for S373–I375 using 2D planes from the 3D spectra. The 2D planes are taken at the $^{15}N$ chemical shift frequency specified for each plot, and the 1D horizonal and vertical slices are the cross sections indicated by the thin, unbroken black line. In Supplementary Figure 5, S374 and S384 are overlapped, but the carbonyl shifts between the two sites are resolved in the 2D planes from the 3D NCACX spectrum shown here. I375 was observed in NCACX and NCOCX spectra as well, but these spectral slices are left off here for sake of space, see supplementary figure 5 for these peaks. All other rigid residues assigned followed the same matching of chemical shift logic, whether assigned by hand or via the MCASSIGN2b program. Contours are drawn at intensity levels increasing by a factor of 1.3 for all spectra.

**Supplementary Table 1. Class C Modeling Statistics.** See methods for additional information.

| | |
|---|---|
| Non-hydrogen atoms | 2196 |
| Number of protein chains | 12 |
| Map CC (mask) | 0.73 |
| RMSZ bonds (Å) | 0.003 |
| RMSZ angles (°) | 0.644 |
| All-atom clash score | 9.96 |
| Ramachandran favored/allowed/outliers (%) | 85.23/14.77/0 |
| Rotamer outliers (%) | 0 |
| CB outliers (%) | 0 |
| Molprobity score | 2.18 |

**Supplementary Table 2. Class B Modeling Statistics.** See methods for additional information.

| | |
|---|---|
| Non-hydrogen atoms | 1062 |
| Number of protein chains | 6 |
| Map CC (mask) | 0.72 |
| RMSZ bonds (Å) | 0.004 |
| RMSZ angles (°) | 0.689 |
| All-atom clash score | 11.14 |
| Ramachandran favored/allowed/outliers (%) | 95.24/4.76/0 |
| Rotamer outliers (%) | 0 |
| CB outliers (%) | 0 |
| Molprobity score | 1.89 |

**Supplementary Table 3. Intra-Class A RMSDs.** RMSD was calculated utilizing ChimeraX (Pettersen, 2021) for residues SY-S373–S392 between the two molecules for all non-hydrogen atoms.

| | Class A Molecule 1 | | | |
|---|---|---|---|---|
| | <u>Backbone</u> | <u>All</u> | <u>Backbone</u><br><u>SY-S373–K376</u> | <u>Backbone</u><br><u>N385–E390</u> |
| Class A Molecule 2 | 1.8 Å | 2.0 Å | 1.0 Å | 1.3 Å |

**Supplementary Table 4. Intra-Class C RMSDs.** RMSD was calculated utilizing ChimeraX (Pettersen, 2021) for residues SY-373–392 between the two molecules for all non-hydrogen atoms.

| | Class C Molecule 1 | |
|---|---|---|
| | <u>Backbone</u> | <u>All</u> |
| Class C Molecule 2 | 1.5 Å | 2.1 Å |

**Supplementary Table 5. Inter-Class RMSDs.** The following RMSDs were calculated utilizing ChimeraX (Pettersen, 2021) for all non-hydrogen atoms. Only residues Y-S373–S392 are used for the alignment as these residues are defined in all models.

| | Class A Molecule 1 | | Class A Molecule 2 | |
|---|---|---|---|---|
| | <u>Backbone</u> | <u>All</u> | <u>Backbone</u> | <u>All</u> |
| Class B | 1.3 Å | 1.8 Å | 0.85 Å | 1.6 Å |
| Class C Molecule 1 | 1.2 Å | 1.8 Å | 1.2 Å | 1.8 Å |
| Class C Molecule 2 | 1.0 Å | 1.7 Å | 1.2 Å | 1.6 Å |

**Supplementary Table 6. Experimental Parameters for Solid State NMR Measurements.**

| **Spectrum** | **Acquisition Parameters**[a,b] | **Processing Parameters**[c] |
|---|---|---|
| 2D $^{13}C$-$^{13}C$ CP-DARR | ns= 16; $\tau_{aq}$ = 10.24 ms ; $\nu_{1H\text{-}carr}$= 3.4 ppm; $\nu_{13C\text{-}carr}$= 99.2 ppm; $\nu_{1H\text{-}CP}$ = 61.2 kHz; $\nu_{13C\text{-}CP}$= 48.5 kHz; $\nu_{1Hdec}$= 83.3 kHz; $\nu_{DARR}$= 12.75 kHz; $\tau_{1H-\pi/2}$=3 µs; $\tau_{13C-\pi/2}$=4 µs; $\tau_{CP}$= 1.0 ms; $\tau_{DARR}$ = 50 ms; $\tau_{1H-dec}$= 6 µs; $\Delta t_1$ = 12 µs; $\tau_{t1}$ = 5.16 ms | GLB$_{t1}$ = 100 Hz<br>GLB$_{t2}$ = 100 Hz |
| 2D $^{13}C$-$^{13}C$ INEPT-TOBSY | ns = 32; $\tau_{aq}$ = 20.48 ms; $\nu_{1H\text{-}carr}$= 3.4 ppm; $\nu_{13C\text{-}carr}$= 41.2 ppm; $\tau_{J/2}$ = 1 ms; $\Delta t_1$ = 24 µs, $\tau_{t1}$ = 5.28 ms; $\tau_{13C-TOBSY}$=38.3 kHz; $\tau_{mix-TOBSY}$=7.54 ms; uses $POST - C9_6^1$ | GLB$_{t1}$ = 100 Hz<br>GLB$_{t2}$ = 100 Hz |
| 2D CP-NCACX | ns= 128; $\tau_{aq}$ = 10.24 ms ; $\nu_{1H\text{-}carr}$= 3.4 ppm; $\nu_{13C\text{-}carr}$= 62.5 ppm; $\nu_{15N\text{-}carr}$= 120.1 ppm; $\nu_{1H\text{-}CP}$ = 48.5 kHz; $\nu_{15N\text{-}CP}$ = 35.7 kHz; $\nu_{15N\text{-}SCP}$ = 34.4 | GLB$_{t1}$ = 50 Hz<br>g3$_{t1}$ = 0.1<br>GLB$_{t2}$ = 100 Hz |

| | | |
|---|---|---|
| | kHz; $\nu_{13CA\text{-}SCP}$= 21.7 kHz; $\nu_{1Hdec}$= 83.3 kHz; $\nu_{DARR}$= 12.75 kHz; $\tau_{1H-\pi/2}$= 3 μs; $\tau_{13C-\pi/2}$= 4 μs; $\tau_{15N-\pi/2}$=6 μs; $\tau_{CP}$= 0.75 ms; $\tau_{SCP}$= 4 ms; $\tau_{DARR}$ = 50 ms; $\tau_{1H-dec}$= 6 μs; $\Delta t_1$ = 108 μs; $\tau_{t1}$ = 9.5 ms | $g3_{t2}$ = 0.1 |
| 2D CP-NCOCX | ns=192; $\tau_{aq}$ = 10.24 ms ; $\nu_{1H\text{-}carr}$= 3.4 ppm; $\nu_{13C\text{-}carr}$= 174.4 ppm; $\nu_{15N\text{-}carr}$= 120.1 ppm; $\nu_{1H\text{-}CP}$ = 48.5 kHz; $\nu_{15N\text{-}CP}$ = 35.7 kHz; $\nu_{15N\text{-}SCP}$ = 35.7 kHz; $\nu_{13CO\text{-}SCP}$=48.5 kHz; $\nu_{1Hdec}$= 83.3 kHz; $\nu_{DARR}$= 12.75 kHz; $\tau_{1H-\pi/2}$= 3 μs; $\tau_{13C-\pi/2}$= 4 μs; $\tau_{15N-\pi/2}$=6 μs; $\tau_{CP}$= 0.75 ms; $\tau_{SCP}$= 4 ms; $\tau_{DARR}$ = 50 ms; $\tau_{1H-dec}$= 6 μs; $\Delta t_1$ = 108 μs; $\tau_{t1}$ = 9.5 ms | $GLB_{t1}$ = 100 Hz<br>$g3_{t1}$ = 0.1<br>$GLB_{t2}$ = 100 Hz<br>$g3_{t2}$ = 0.1 |
| 3D CP-NCACX | ns= 32; $\tau_{aq}$ = 15.36 ms ; $\nu_{1H\text{-}carr}$= 3.4 ppm; $\nu_{13C\text{-}carr}$= 62.5 ppm; $\nu_{15N\text{-}carr}$= 120.1 ppm; $\nu_{1H\text{-}CP}$ = 48.5 kHz; $\nu_{15N\text{-}CP}$ = 35.7 kHz; $\nu_{15N\text{-}SCP}$ = 34.4 kHz; $\nu_{13CA\text{-}SCP}$= 21.7 kHz; $\nu_{1Hdec}$= 83.3 kHz; $\nu_{DARR}$= 12.75 kHz; $\tau_{1H-\pi/2}$= 3 μs; $\tau_{13C-\pi/2}$= 4 μs; $\tau_{15N-\pi/2}$=6 μs; $\tau_{CP}$= 0.75 ms; $\tau_{SCP}$= 4 ms; $\tau_{DARR}$ = 50 ms; $\tau_{1H-dec}$= 6 μs; $\Delta t_1$ = 180 μs; $\tau_{t1}$ = 7.56 ms; $\Delta t_2$ = 132.0 μs; $\tau_{t2}$ = 3.3 ms | $GLB_{t1}$ = 60 Hz<br>$GLB_{t2}$ = 80 Hz<br>$GLB_{t3}$ = 100 Hz |
| 3D CP-NCOCX | ns= 40; $\tau_{aq}$ = 15.36 ms ; $\nu_{1H\text{-}carr}$= 3.4 ppm; $\nu_{13C\text{-}carr}$= 174.4 ppm; $\nu_{15N\text{-}carr}$= 120.1 ppm; $\nu_{1H\text{-}CP}$ = 48.5 kHz; $\nu_{15N\text{-}CP}$ = 35.7 kHz; $\nu_{15N\text{-}SCP}$ = 35.7 kHz; $\nu_{13CO\text{-}SCP}$=48.5 kHz; $\nu_{1Hdec}$= 83.3 kHz; $\nu_{DARR}$= 12.75 kHz; $\tau_{1H-\pi/2}$= 3 μs; $\tau_{13C-\pi/2}$= 4 μs; $\tau_{15N-\pi/2}$=6 μs; $\tau_{CP}$= 0.75 ms; $\tau_{SCP}$= 4 ms; $\tau_{DARR}$ = 50 ms; $\tau_{1H-dec}$= 6 μs; $\Delta t_1$ = 180 μs; $\tau_{t1}$ = 7.56 ms; $\Delta t_2$ = 132.0 μs; $\tau_{t2}$ = 3.3 ms | $GLB_{t1}$ = 100 Hz<br>$GLB_{t2}$ = 80 Hz<br>$GLB_{t3}$ = 100 Hz |
| 3D CP-CANCO | ns= 16; $\tau_{aq}$ = 10.24 ms ; $\nu_{1H\text{-}carr}$= 3.4 ppm; $\nu_{13C\text{-}carr}$= 62.5 ppm; $\nu_{15N\text{-}carr}$= 120.1 ppm; $\nu_{1H\text{-}CP}$ = 48.5 kHz; $\nu_{15N\text{-}CP}$ = 35.7 kHz $\nu_{15N\text{-}SCP}$ = 34.4 kHz; $\nu_{13CA\text{-}SCP}$= 21.7 kHz; $\nu_{13CO-SCP-eff}$= 47.2 kHz; $\nu_{1Hdec}$= 83.3 kHz; $\nu_{DARR}$= 12.75 kHz; $\tau_{1H-\pi/2}$= 3 μs; $\tau_{13C-\pi/2}$= 4 μs; $\tau_{15N-\pi/2}$=6 μs; $\tau_{CP}$= 0.75 ms; $\tau_{SCP}$= 4 ms; $\tau_{DARR}$ = 50 ms; $\tau_{1H-dec}$= 6 μs; $\Delta t_2$ = 180 μs; $\tau_{t2}$ = 7.56 ms; $\Delta t_1$ = 60 μs; $\tau_{t1}$ = 3.3 ms | $GLB_{t1}$ = 100 Hz<br>$GLB_{t2}$ = 100 Hz<br>$GLB_{t3}$ = 50 Hz<br>$g3_{t3}$ = 0.1 |

[a]All of the $^{1}$H-X CP transfers use a 20% ramp on the $^{1}$H channel. All $^{15}$N-$^{13}$C Specific-CP transfers used a 5% ramp on the $^{15}$N channel.

[b]ns is the number of scans averaged; $\tau_{aq}$ is the total acquisition time in the directly detected dimension; $\nu_{X\text{-}carr}$ is the center frequency; $\nu_{X\text{-}CP}$ is the nutation frequency of the applied power during the CP step; $\nu_{X\text{-}SCP}$ is the nutation frequency from the applied power during the specific CP step; $\nu_{1Hdec}$ is the decoupling pulse power for $^{1}$H; $\nu_{DARR}$ is the $^{13}$C – $^{13}$C mixing pulse power on $^{1}$H; $\tau_{X-\pi/2}$ is the nuclei specified 90° pulse length; $\tau_{13C-TOBSY}$ is the $^{13}$C TOBSY pulse length; $\tau_{CP}$ is the contact time for CP; $\tau_{SCP}$ is the contact time for specific CP; $\tau_{DARR}$ is the $^{13}$C-$^{13}$C DARR mixing time; $\tau_{mix-TOBSY}$ is the total TOBSY mixing time; $\tau_{1H-dec}$ is

the SPINAL64 π pulse length; $\Delta t_x$is the increment in the specified indirect time dimension; $\tau_{tx}$ is the total acquisition time for the specified dimension, $\tau_{J/2}$is the ½ echo delay; $\nu_{13CO-SCP-eff}$ is the effective SCP pulse to $^{13}C$ carbonyl carbon from $^{15}N$.

[c] Spectra are processed in NMRPipe. Gaussian line broadening was applied according to the parameters specified. Zero filling was applied twice before applying each Fourier transform.

**Supplementary Table 7. Residues Assigned from the Cross Polarization-Based Spectra**

| Residue | Type | N | CA | CO | CB | CG | CD | S/N[a] |
|---|---|---|---|---|---|---|---|---|
| | **S** | 118.0 | 55.6 | 172.9 | 66.3 | | | |
| | **Y** | 116.6 | 54.1 | 176.4 | 41.0 | | | 11* |
| **373** | **S** | 115.8 | 56.5 | 173.1 | 67.4 | | | 10 |
| **374** | **S** | 114.6 | 56.1 | 173.6 | 64.1 | | | |
| **375** | **I** | 127.9 | 60.5 | 174.5 | 41.9 | 27.5 | 17.2 | 23 |
| **376** | **K** | 129.7 | 55.0 | 174.0 | 35.7 | 25.6 | 29.9 | 31 |
| **377** | **L** | 127.2 | 53.8 | 175.9 | 43.8 | 29.1 | 26.6 | 33 |
| **378** | **S** | 120.0 | 55.6 | 173.0 | 65.4 | | | 26 |
| **379** | **N** | 118.1 | 53.9 | 172.9 | 36.8 | 178.5 | | 20 |
| **380** | **N** | 114.4 | 52.9 | 173.9 | 39.3 | 176.8 | | 29* |
| **381** | **N** | 121.4 | 52.6 | 174.9 | 42.5 | | | 25 |
| **382** | **N** | 112.3 | 54.1 | 174.0 | 36.4 | 178.4 | | 18 |
| **383** | **N** | 114.2 | 54.1 | 174.1 | 42.2 | 177.7 | | 32 |
| **384** | **S** | 114.5 | 56.0 | 175.5 | 64.1 | | | 32 |
| **385** | **N** | 128.6 | 54.3 | 173.8 | 37.5 | 178.1 | | 32 |
| **386** | **S** | 111.0 | 58.8 | 171.3 | 69.2 | | | 59 |
| **387** | **N** | 115.1 | 52.7 | 172.1 | 36.7 | 176.3 | | 29 |
| **388** | **N** | 113.4 | 53.1 | 174.9 | 40.1 | | | 47 |
| **389** | **I** | 120.0 | 60.4 | 174.2 | 41.3 | 27.1 | 17.0 | 19** |
| **390** | **E** | 127.2 | 54.3 | 174.3 | 32.9 | 36.1 | 183.5 | 33 |
| **391** | **I** | 125.0 | 59.8 | 174.5 | 41.1 | 27.0 | 16.8 | 28 |
| **392** | **S** | 120.4 | 55.8 | 173.9 | 65.5 | | | 20* |
| **393** | **K** | 128.2 | | | | | | |
| **401** | **D** | | 53.0 | 175.7 | 39.3 | 177.1 | | |
| **402** | **I** | 122.5 | 58.8 | 174.6 | 40.4 | 27.0 | 16.9 | 15 |
| **403** | **G** | 115.9 | 43.3 | 172.1 | | | | 11 |
| **404** | **G** | 106.2 | 44.5 | 175.2 | | | | 11 |
| **405** | **T** | 120.3 | 60.4 | 172.5 | 71.9 | 21.2 | | 19** |
| **406** | **N** | 121.3 | 53.7 | 173.2 | 36.5 | | | 17 |

| [a]S/N is the value calculated by NMRFAM-SPARKY for the CANCO peaks. * represents overlapped peaks, and ** is the resonance that is used twice in the assignments for the CANCO peak (but is unique in the NCACX and NCOCX spectra). |
|---|

**Supplementary Table 8. Residues Assigned from the J-Based INEPT-TOBSY Spectrum**

| Residue | Type | CA | CB | CG | CD |
|---|---|---|---|---|---|
| **410** | **A** | **52.2** | **19.0** | | |
| **411** | **S** | **58.3** | **63.5** | | |
| **412** | **R** | 56.1 | 30.7 | 27.0 | 43.1 |
| **414** | **I** | 60.9 | 38.5 | 27.1 | 17.3 |
| **415** | **A** | **52.2** | **19.0** | | |
| **416** | **S** | **58.3** | **63.5** | | |
| **417** | **A** | **52.2** | **19.0** | | |
| **418** | **A** | **52.2** | **19.0** | | |
| **419** | **V** | 62.3 | 32.6 | 20.5 | |
| **420** | **G** | 45.0 | | | |
| **421** | **E** | **56.4** | **30.0** | **36.0** | |
| **422** | **E** | **56.4** | **30.0** | **36.0** | |
| **424** | **S** | **58.3** | **63.5** | | |
| **426** | **L** | 55.2 | 42.1 | 26.8 | 24.8 |
| **427** | **S** | **58.3** | **63.5** | | |
| **428** | **S** | **58.3** | **63.5** | | |
| **430** | **S** | **58.3** | **63.5** | | |
| **432** | **E** | **56.4** | **30.0** | **36.0** | |
| **434** | **N** | **53.1** | **38.7** | | |
| **435** | **N** | **53.1** | **38.7** | | |
| **436** | **N** | 51.1 | 38.6 | | |
| **437** | **P** | 63.5 | 31.9 | 26.9 | 50.4 |
| **438** | **N** | **53.1** | **38.7** | | |
| **439** | **N** | **53.1** | **38.7** | | |
| **440** | **D** | 54.4 | 40.9 | | |
| RED represents the tentative assignments made for multiple residues via the process detailed in the methods section. | | | | | |

**Supplementary Table 9. Unassigned Resonances from the Cross Polarization-Based NCACX Experiment**

| N | CA | CO | CB | CG | CD | Residue Type |
|---|---|---|---|---|---|---|
| 127.9 | 60.4 | 174.4 | 42.4 | 27.7 | 18.5 | **I*** |
| 125.1 | 56.0 | 172.3 | 64.2 | | | **S** |
| 124.8 | 56.4 | 170.8 | 64.9 | | | **S** |
| 126.0 | 51.2 | 172.7 | 23.1 | | | **A** |
| 114.3 | 54.0 | 174.0 | 36.5 | | | **RDNCEQHKMFWY** |

| | | | | | | |
|---|---|---|---|---|---|---|
| 115.1 | 52.7 | 174.0 | 35.9 | | | **RNCEQHKMFWY** |
| 122.9 | 61.6 | 173.0 | 70.8 | 21.2 | | **T** |
| 117.3 | 56.2 | 173.1 | 65.3 | | | **S** |
| 123.7 | 50.9 | 174.1 | 21.3 | | | **A** |
| 124.9 | 53.8 | 176.2 | 29.0 | 24.3 | 43.8 | **RCEQHKMW** |
| 128.9 | 54.2 | 174.5 | 32.9 | | | **RCEQHIKMWY** |

* The I peak could be an alternative assignment to I375, where the chemical shift difference is ~0.4 ppm for the CB residue and ~1.3 ppm for the CD residue relative to the assigned chemical shifts for I375 in Supplemental Table 7, while the other I375 atoms have the nearly identical chemical shifts. This might indicate alternative side chain packing arrangement for I375, although this is not able to be observed in the reconstructed cryo-EM map.

**Supplementary Table 10. Unassigned Resonances from INEPT-TOBSY J-Based Spectrum**

| CA | CB | CG | Residue Type |
|---|---|---|---|
| 61.8 | 69.5 | 21.4 | **T** |
| 63.1 | 70.6 | 21.8 | **T** |
| | 41.7 | 138.2 | **?** |

**Supplementary Table 11. NCACX/DARR Signal Table For MCASSIGN2b.**

| Observed Chemical Shift (ppm) | | | | | | Chemical Shift Uncertainty (ppm) | | | | | | | |
|---|---|---|---|---|---|---|---|---|---|---|---|---|---|
| $^{15}$N | $^{13}$CA | $^{13}$CO | $^{13}$CB | $^{13}$CG | $^{13}$CD /CG2 | $^{15}$N | $^{13}$CA | $^{13}$CO | $^{13}$CB | $^{13}$CG | $^{13}$CD /CG2 | MP[a] | Residue Type |
| 106.38 | 44.68 | 175.18 | 1111.1 | 1111.1 | 1111.1 | 0.5 | 0.5 | 0.5 | 0.5 | 0.5 | 0.5 | 1 | G |
| 111.03 | 58.90 | 171.48 | 69.18 | 1111.10 | 1111.10 | 0.5 | 0.5 | 0.5 | 0.5 | 0.5 | 0.5 | 1 | ST |
| 112.03 | 54.22 | 173.96 | 36.47 | 178.37 | 1111.10 | 0.5 | 0.5 | 0.5 | 0.5 | 0.5 | 0.5 | 1 | RDNCEHIK Y |
| 114.08 | 53.04 | 174.07 | 39.29 | 176.58 | 1111.10 | 0.5 | 0.5 | 0.5 | 0.5 | 0.5 | 0.5 | 1 | DNCLY |
| 114.66 | 56.04 | 175.59 | 63.94 | 1111.10 | 1111.10 | 0.5 | 0.5 | 0.5 | 0.5 | 0.5 | 0.5 | 1 | ST |
| 114.44 | 54.10 | 174.07 | 42.23 | 177.60 | 1111.10 | 0.5 | 0.5 | 0.5 | 0.5 | 0.5 | 0.5 | 1 | DNCILY |
| 115.11 | 52.76 | 172.14 | 36.67 | 176.37 | 1111.10 | 0.5 | 0.5 | 2.0 | 0.5 | 2.0 | 0.5 | 1 | RDNCEHKY |
| 128.79 | 54.40 | 173.88 | 37.51 | 178.14 | 1111.10 | 0.5 | 0.5 | 0.5 | 0.5 | 0.5 | 0.5 | 1 | DNCHILKY |
| 121.64 | 52.55 | 174.95 | 42.53 | 1111.10 | 1111.10 | 0.5 | 0.5 | 0.5 | 0.5 | 0.5 | 0.5 | 1 | DNCLY |
| 118.22 | 54.04 | 172.99 | 36.96 | 178.48 | 1111.10 | 0.5 | 0.5 | 0.5 | 0.5 | 0.5 | 0.5 | 1 | RDNCEHIK Y |
| 115.96 | 56.53 | 173.20 | 67.42 | 1111.10 | 1111.10 | 0.5 | 0.5 | 0.5 | 0.5 | 0.5 | 0.5 | 1 | ST |
| 119.95 | 60.57 | 174.43 | 41.13 | 27.07 | 17.03 | 0.5 | 0.5 | 0.5 | 0.5 | 0.5 | 0.5 | 1 | CILY |
| 116.04 | 43.24 | 172.10 | 1111.10 | 1111.10 | 1111.10 | 0.5 | 0.5 | 0.5 | 0.5 | 0.5 | 0.5 | 1 | G |
| 125.14 | 59.84 | 174.34 | 41.03 | 26.97 | 16.87 | 0.5 | 0.5 | 0.5 | 0.5 | 0.5 | 0.5 | 1 | CILY |
| 127.94 | 60.44 | 174.44 | 42.40 | 27.69 | 18.50 | 0.5 | 0.5 | 0.5 | 0.5 | 0.5 | 0.5 | 1 | CILY |
| 128.01 | 60.51 | 174.44 | 41.86 | 27.37 | 17.14 | 0.5 | 0.5 | 0.5 | 0.5 | 0.5 | 0.5 | 1 | CILY |
| 125.14 | 56.00 | 172.33 | 64.20 | 1111.10 | 1111.10 | 0.5 | 0.5 | 0.5 | 0.5 | 0.5 | 0.5 | 1 | ST |
| 124.84 | 56.42 | 170.80 | 64.93 | 1111.10 | 1111.10 | 0.5 | 0.5 | 0.5 | 0.5 | 0.5 | 0.5 | 1 | ST |

| | | | | | | | | | | | | | |
|---|---|---|---|---|---|---|---|---|---|---|---|---|---|
| 127.03 | 54.28 | 174.40 | 32.92 | 36.02 | 183.44 | 0.5 | 0.5 | 0.5 | 0.5 | 0.5 | 0.5 | 1 | RCEHIKY |
| 122.67 | 58.71 | 174.19 | 40.35 | 27.04 | 16.94 | 0.5 | 0.5 | 0.5 | 0.5 | 0.5 | 0.5 | 1 | DNCILY |
| 119.91 | 55.76 | 173.06 | 65.33 | 1111.10 | 1111.10 | 0.5 | 0.5 | 0.5 | 0.5 | 0.5 | 0.5 | 1 | ST |
| 120.52 | 55.90 | 173.89 | 65.77 | 1111.10 | 1111.10 | 0.5 | 0.5 | 0.5 | 0.5 | 0.5 | 0.5 | 1 | ST |
| 121.46 | 53.67 | 173.15 | 36.52 | 1111.10 | 1111.10 | 0.5 | 0.5 | 0.5 | 0.5 | 0.5 | 0.5 | 1 | RDNCEHKY |
| 129.65 | 55.19 | 173.93 | 35.52 | 25.60 | 29.90 | 0.5 | 0.5 | 0.5 | 0.5 | 0.5 | 0.5 | 1 | RNCEHIKY |
| 125.99 | 51.24 | 172.73 | 23.07 | 1111.10 | 1111.10 | 0.5 | 0.5 | 0.5 | 0.5 | 0.5 | 0.5 | 1 | AC |
| 120.49 | 60.43 | 172.61 | 72.00 | 21.24 | 1111.10 | 0.5 | 0.5 | 0.5 | 0.5 | 0.5 | 0.5 | 1 | T |
| 122.23 | 62.12 | 173.78 | 32.57 | 21.00 | 1111.10 | 0.5 | 0.5 | 0.5 | 0.5 | 0.5 | 0.5 | 1 | RCEHIV |
| 114.25 | 53.95 | 174.02 | 36.48 | 1111.10 | 1111.10 | 0.5 | 0.5 | 0.5 | 0.5 | 0.5 | 0.5 | 1 | RDNCEHKY |
| 114.59 | 56.01 | 173.76 | 64.15 | 1111.10 | 1111.10 | 0.5 | 0.5 | 0.5 | 0.5 | 0.5 | 0.5 | 1 | ST |
| 127.11 | 53.66 | 176.10 | 43.81 | 29.20 | 26.62 | 0.5 | 0.5 | 0.5 | 0.5 | 0.5 | 0.5 | 1 | DNCLY |
| 115.10 | 52.75 | 173.98 | 35.89 | 1111.10 | 1111.10 | 0.5 | 0.5 | 2.0 | 0.5 | 0.5 | 0.5 | 1 | RNCEHKY |
| 122.92 | 61.57 | 173.00 | 70.78 | 21.17 | 1111.10 | 0.5 | 0.5 | 0.5 | 0.5 | 0.5 | 0.5 | 1 | T |
| 118.00 | 55.50 | 172.95 | 66.29 | 1111.10 | 1111.10 | 0.5 | 0.5 | 1.0 | 0.5 | 0.5 | 0.5 | 1 | S |
| 117.29 | 56.24 | 173.09 | 65.28 | 1111.10 | 1111.10 | 0.5 | 1.0 | 1.0 | 0.5 | 0.5 | 0.5 | 1 | ST |
| 113.42 | 53.18 | 174.79 | 39.99 | 1111.10 | 1111.10 | 0.5 | 0.5 | 0.5 | 0.5 | 0.5 | 0.5 | 1 | DNCLY |
| 116.79 | 54.08 | 176.56 | 41.12 | 1111.10 | 1111.10 | 0.5 | 0.5 | 0.5 | 0.5 | 0.5 | 0.5 | 1 | DNCILY |
| 123.73 | 50.91 | 174.12 | 21.26 | 1111.10 | 1111.10 | 0.5 | 0.5 | 0.5 | 0.5 | 0.5 | 0.5 | 1 | A |
| 124.90 | 53.80 | 176.20 | 29.02 | 24.31 | 43.80 | 0.5 | 0.5 | 0.5 | 0.5 | 0.5 | 0.5 | 1 | RCEHK |
| 128.90 | 54.23 | 174.50 | 32.86 | 1111.10 | 1111.10 | 0.5 | 0.5 | 0.5 | 0.5 | 0.5 | 0.5 | 1 | RCEHIKY |

[a]multiplicity (MP), the number of times a signal can be assigned

#### Supplementary Table 12. NCOCX Signal Table For MCASSIGN2b.

| Observed Chemical Shift (ppm) | | | | | | Chemical Shift Uncertainty (ppm) | | | | | | | |
|---|---|---|---|---|---|---|---|---|---|---|---|---|---|
| $^{15}$N | $^{13}$CA | $^{13}$CO | $^{13}$CB | $^{13}$CG | $^{13}$CD /CG2 | $^{15}$N | $^{13}$CA | $^{13}$CO | $^{13}$CB | $^{13}$CG | $^{13}$CD /CG2 | MP[a] | Residue Type |
| 106.18 | 43.36 | 172.11 | 1111.10 | 1111.10 | 1111.10 | 0.5 | 0.5 | 0.5 | 0.5 | 0.5 | 0.5 | 1 | G |
| 111.08 | 54.36 | 173.73 | 37.50 | 178.16 | 1111.10 | 0.5 | 0.5 | 0.5 | 0.5 | 0.5 | 0.5 | 1 | DNCHILKY |
| 112.25 | 53.21 | 176.68 | 39.13 | 1111.10 | 1111.10 | 0.5 | 0.5 | 0.5 | 0.5 | 0.5 | 0.5 | 1 | DNCLY |
| 112.41 | 52.82 | 174.77 | 42.43 | 1111.10 | 1111.10 | 0.5 | 0.5 | 0.5 | 0.5 | 0.5 | 0.5 | 1 | DNCLY |
| 113.38 | 52.78 | 172.12 | 36.75 | 176.30 | 1111.10 | 0.5 | 0.5 | 0.5 | 0.5 | 0.5 | 0.5 | 1 | RDNCEHKY |
| 114.38 | 54.15 | 174.09 | 42.19 | 177.74 | 1111.10 | 0.5 | 0.5 | 0.5 | 0.5 | 0.5 | 0.5 | 1 | DNCILY |
| 113.96 | 54.01 | 174.08 | 36.32 | 1111.10 | 1111.10 | 0.5 | 0.5 | 0.5 | 0.5 | 0.5 | 0.5 | 1 | RDNCEHIKY |
| 115.20 | 58.90 | 171.33 | 69.21 | 1111.10 | 1111.10 | 0.5 | 0.5 | 0.5 | 0.5 | 0.5 | 0.5 | 1 | ST |
| 114.72 | 53.90 | 173.04 | 36.66 | 178.55 | 1111.10 | 0.5 | 0.5 | 0.5 | 0.5 | 0.5 | 0.5 | 1 | RDNCEHKY |
| 118.10 | 55.55 | 173.04 | 65.39 | 1111.10 | 1111.10 | 0.5 | 0.5 | 0.5 | 0.5 | 0.5 | 0.5 | 1 | ST |
| 120.31 | 59.91 | 174.75 | 41.07 | 27.02 | 16.81 | 0.5 | 0.5 | 0.5 | 0.5 | 0.5 | 0.5 | 1 | CILY |
| 120.05 | 56.00 | 173.35 | 65.79 | 1111.10 | 1111.10 | 0.5 | 0.5 | 0.5 | 0.5 | 0.5 | 0.5 | 1 | ST |
| 120.63 | 55.82 | 172.45 | 64.10 | 1111.10 | 1111.10 | 0.5 | 0.5 | 0.5 | 0.5 | 0.5 | 0.5 | 1 | ST |
| 120.87 | 60.87 | 172.62 | 42.25 | 27.13 | 1111.10 | 0.5 | 0.5 | 0.5 | 0.5 | 0.5 | 0.5 | 1 | CILY |

| | | | | | | | | | | | | | |
|---|---|---|---|---|---|---|---|---|---|---|---|---|---|
| 125.00 | 54.48 | 174.25 | 32.88 | 36.14 | 183.47 | 0.5 | 0.5 | 0.5 | 0.5 | 0.5 | 0.5 | 1 | RCEHIKY |
| 124.73 | 55.83 | 174.01 | 1111.10 | 1111.10 | 1111.10 | 0.5 | 0.5 | 0.5 | 0.5 | 0.5 | 0.5 | 1 | RDNCEHILKSTY |
| 129.78 | 60.70 | 174.54 | 42.02 | 27.69 | 17.27 | 0.5 | 0.5 | 0.5 | 0.5 | 0.5 | 0.5 | 1 | CILY |
| 128.49 | 55.96 | 175.45 | 64.16 | 1111.10 | 1111.10 | 0.5 | 0.5 | 0.5 | 0.5 | 0.5 | 0.5 | 1 | ST |
| 127.68 | 55.44 | 173.70 | 1111.10 | 1111.10 | 1111.10 | 0.5 | 0.5 | 0.5 | 0.5 | 0.5 | 0.5 | 1 | ARDNCEHILKSY |
| 127.84 | 56.25 | 173.42 | 64.05 | 1111.10 | 1111.10 | 0.5 | 0.5 | 0.5 | 0.5 | 0.5 | 0.5 | 1 | ST |
| 128.15 | 55.53 | 173.91 | 65.15 | 1111.10 | 1111.10 | 0.5 | 0.5 | 0.5 | 0.5 | 0.5 | 0.5 | 1 | ST |
| 125.45 | 56.29 | 173.20 | 64.89 | 1111.10 | 1111.10 | 0.5 | 0.5 | 0.5 | 0.5 | 0.5 | 0.5 | 1 | ST |
| 123.35 | 58.12 | 172.40 | 1111.10 | 1111.10 | 1111.10 | 0.5 | 0.5 | 0.5 | 0.5 | 0.5 | 0.5 | 1 | RDNCEHILKPSTYV |
| 133.29 | 55.26 | 171.93 | 1111.10 | 1111.10 | 1111.10 | 0.5 | 0.5 | 0.5 | 0.5 | 0.5 | 0.5 | 1 | ARDNCEHILKSY |
| 123.10 | 55.80 | 173.77 | 66.14 | 1111.10 | 1111.10 | 0.5 | 0.5 | 0.5 | 0.5 | 0.5 | 0.5 | 1 | ST |
| 120.17 | 44.20 | 175.48 | 1111.10 | 1111.10 | 1111.10 | 0.5 | 0.5 | 0.5 | 0.5 | 0.5 | 0.5 | 1 | G |
| 122.38 | 53.01 | 175.72 | 39.28 | 177.10 | 1111.10 | 0.5 | 0.5 | 0.5 | 0.5 | 0.5 | 0.5 | 1 | DNCLY |
| 121.22 | 52.86 | 173.85 | 39.25 | 177.01 | 1111.10 | 0.5 | 0.5 | 0.5 | 0.5 | 0.5 | 0.5 | 1 | DNCLY |
| 121.76 | 52.95 | 175.23 | 39.85 | 177.13 | 1111.10 | 0.5 | 0.5 | 0.5 | 0.5 | 0.5 | 0.5 | 1 | DNCLY |
| 121.24 | 60.42 | 172.50 | 71.87 | 21.25 | 1111.10 | 0.5 | 0.5 | 0.5 | 0.5 | 0.5 | 0.5 | 1 | T |
| 127.31 | 60.32 | 174.12 | 41.41 | 27.06 | 16.95 | 0.5 | 0.5 | 0.5 | 0.5 | 0.5 | 0.5 | 1 | CILY |
| 113.38 | 52.78 | 172.13 | 35.97 | 1111.10 | 1111.10 | 0.5 | 0.5 | 0.5 | 0.5 | 0.5 | 0.5 | 1 | RNCEHKY |
| 114.61 | 56.73 | 172.98 | 65.48 | 1111.10 | 1111.10 | 0.5 | 0.5 | 0.5 | 0.5 | 0.5 | 0.5 | 1 | ST |
| 114.61 | 56.75 | 173.06 | 67.39 | 1111.10 | 1111.10 | 0.5 | 0.5 | 0.5 | 0.5 | 0.5 | 0.5 | 1 | ST |
| 124.71 | 57.18 | 175.14 | 42.18 | 1111.10 | 1111.10 | 0.5 | 0.5 | 0.5 | 0.5 | 0.5 | 0.5 | 1 | DNCILY |
| 120.01 | 53.82 | 175.75 | 43.76 | 29.00 | 1111.10 | 0.5 | 0.5 | 0.5 | 0.5 | 0.5 | 0.5 | 1 | DNCLY |
| 115.82 | 59.07 | 174.73 | 1111.10 | 27.04 | 16.79 | 0.5 | 0.5 | 0.5 | 0.5 | 0.5 | 0.5 | 1 | RDNCEHILKPSTYV |
| 127.43 | 55.00 | 174.05 | 35.86 | 1111.10 | 1111.10 | 0.5 | 0.5 | 0.5 | 0.5 | 0.5 | 0.5 | 1 | RNCEHIKY |
| 119.99 | 53.22 | 174.82 | 40.26 | 1111.10 | 1111.10 | 0.5 | 0.5 | 0.5 | 0.5 | 0.5 | 0.5 | 1 | DNCLY |
| [a]multiplicity (MP), the number of times a signal can be assigned | | | | | | | | | | | | | |

#### Supplementary Table 13. CANCO Signal Table For MCASSIGN2b.

| Observed Chemical Shift (ppm) | | | Chemical Shift Uncertainty (ppm) | | | | |
|---|---|---|---|---|---|---|---|
| $^{15}$N | $^{13}$CA | $^{13}$CO | $^{15}$N | $^{13}$CA | $^{13}$CO | MP[a] | Residue Type |
| 106.05 | 44.49 | 172.01 | 0.7 | 0.7 | 0.7 | 1 | G |
| 110.88 | 58.68 | 173.64 | 0.7 | 0.7 | 0.7 | 1 | RDNCEHILKSTYV |
| 112.59 | 53.96 | 174.91 | 0.7 | 0.7 | 0.7 | 1 | ARDNCEHLKSY |
| 113.31 | 52.99 | 172.08 | 0.7 | 0.7 | 0.7 | 1 | ARDNCEHLKSY |
| 114.45 | 55.96 | 174.06 | 0.7 | 0.7 | 0.7 | 1 | RDNCEHILKSTY |
| 114.25 | 53.97 | 174.05 | 0.7 | 0.7 | 0.7 | 1 | ARDNCEHLKSY |
| 114.36 | 52.94 | 172.65 | 0.7 | 0.7 | 0.7 | 1 | ARDNCEHLKSY |
| 114.93 | 52.60 | 171.20 | 0.7 | 0.7 | 0.7 | 1 | ARDNCEHLKSY |
| 128.62 | 54.19 | 175.45 | 0.7 | 0.7 | 0.7 | 1 | ARDNCEHILKSY |
| 121.23 | 52.46 | 173.82 | 0.7 | 0.7 | 0.7 | 1 | ARDNCEHLKSY |
| 118.01 | 53.90 | 172.94 | 0.7 | 0.7 | 0.7 | 1 | ARDNCEHLKSY |

| | | | | | | | |
|---|---|---|---|---|---|---|---|
| 115.87 | 56.31 | 176.02 | 0.7 | 0.7 | 0.7 | 1 | RDNCEHILKSTYV |
| 120.11 | 60.45 | 174.95 | 0.7 | 0.7 | 0.7 | 2 | RCEHILKSTYV |
| 115.84 | 43.28 | 175.00 | 0.7 | 0.7 | 0.7 | 1 | G |
| 124.91 | 59.65 | 174.30 | 0.7 | 0.7 | 0.7 | 1 | RCEHILKSTYV |
| 127.91 | 60.36 | 173.70 | 0.7 | 0.7 | 0.7 | 1 | RCEHILKSTYV |
| 127.91 | 60.36 | 173.70 | 0.7 | 0.7 | 0.7 | 1 | RCEHILKSTYV |
| 125.08 | 55.76 | 174.83 | 0.7 | 0.7 | 0.7 | 1 | RDNCEHILKSTY |
| 124.58 | 56.30 | 175.61 | 0.7 | 0.7 | 0.7 | 1 | RDNCEHILKSTYV |
| 127.14 | 54.03 | 174.08 | 0.7 | 0.7 | 0.7 | 1 | ARDNCEHILKSY |
| 122.52 | 58.58 | 175.98 | 0.7 | 0.7 | 0.7 | 1 | RDNCEHILKSTYV |
| 119.97 | 55.40 | 175.82 | 0.7 | 0.7 | 0.7 | 1 | ARDNCEHILKSY |
| 120.71 | 55.76 | 172.29 | 0.7 | 0.7 | 0.7 | 1 | RDNCEHILKSTY |
| 121.25 | 53.55 | 172.34 | 0.7 | 0.7 | 0.7 | 1 | ARDNCEHLKSY |
| 129.61 | 54.95 | 174.58 | 0.7 | 0.7 | 0.7 | 1 | ARDNCEHILKSY |
| 120.24 | 55.83 | 174.50 | 0.7 | 0.7 | 0.7 | 1 | RDNCEHILKSTY |
| 120.22 | 55.98 | 173.92 | 0.7 | 0.7 | 0.7 | 1 | RDNCEHILKSTY |
| 120.10 | 54.93 | 170.49 | 0.7 | 0.7 | 0.7 | 1 | ARDNCEHILKSY |
| 121.83 | 52.27 | 175.27 | 0.7 | 0.7 | 0.7 | 1 | ARDNCEHLKSY |
| 127.92 | 55.10 | 173.10 | 0.7 | 0.7 | 0.7 | 1 | ARDNCEHILKSY |
| 116.80 | 53.90 | 172.96 | 0.7 | 0.7 | 0.7 | 1 | ARDNCEHLKSY |
| 123.89 | 50.24 | 174.51 | 0.7 | 0.7 | 0.7 | 1 | ADNCEHLK |
| 128.63 | 54.21 | 173.97 | 0.7 | 0.7 | 0.7 | 1 | ARDNCEHILKSY |

[a]multiplicity (MP), the number of times a signal can be assigned
